# Single-cell analysis of bidirectional reprogramming between early embryonic states reveals mechanisms of differential lineage plasticities

**DOI:** 10.1101/2023.03.28.534648

**Authors:** Vidur Garg, Yang Yang, Sonja Nowotschin, Manu Setty, Ying-Yi Kuo, Roshan Sharma, Alexander Polyzos, Eralda Salataj, Dylan Murphy, Amy Jang, Dana Pe’er, Effie Apostolou, Anna-Katerina Hadjantonakis

**Affiliations:** Developmental Biology Program, Sloan Kettering Institute, Memorial Sloan Kettering Cancer Center, New York, NY 10065, USA; Biochemistry, Cell and Molecular Biology Program, Weill Cornell Graduate School of Medical Sciences, New York, NY 10021, USA; Computational and Systems Biology Program, Sloan Kettering Institute, Memorial Sloan Kettering Cancer Center, New York, NY 10065, USA; Joan & Sanford I. Weill Department of Medicine, Sandra and Edward Meyer Cancer Center, Weill Cornell Medicine, New York, NY 10021, USA; Howard Hughes Medical Institute, New York, NY 10065, USA

**Keywords:** epiblast, primitive endoderm, ES cells, XEN cells, pluripotency, extra-embryonic endoderm, reprogramming, lineage plasticity, single-cell analysis, blastocyst

## Abstract

Two distinct fates, pluripotent epiblast (EPI) and primitive (extra-embryonic) endoderm (PrE), arise from common progenitor cells, the inner cell mass (ICM), in mammalian embryos. To study how these sister identities are forged, we leveraged embryonic (ES) and e*X*traembryonic *EN*doderm (XEN) stem cells – *in vitro* counterparts of the EPI and PrE. Bidirectional reprogramming between ES and XEN coupled with single-cell RNA and ATAC-seq analyses uncovered distinct rates, efficiencies and trajectories of state conversions, identifying drivers and roadblocks of reciprocal conversions. While GATA4-mediated ES-to-iXEN conversion was rapid and nearly deterministic, OCT4, KLF4 and SOX2-induced XEN-to-iPS reprogramming progressed with diminished efficiency and kinetics. The dominant PrE transcriptional program, safeguarded by *Gata4*, and globally elevated chromatin accessibility of EPI underscored the differential plasticities of the two states. Mapping *in vitro* trajectories to embryos revealed reprogramming in either direction tracked along, and toggled between, EPI and PrE *in vivo* states without transitioning through the ICM.

## INTRODUCTION

In mammals, the pluripotent epiblast (EPI), one of the first lineages specified during embryo development, emerges contemporaneously with its sister lineage, the extra-embryonic (primitive) endoderm (PrE), from a common progenitor population, the inner cell mass (ICM) (Chazaud and Yamanaka, 2016; Schrode et al., 2013). The EPI and PrE lineages are distinct, with the EPI giving rise to the embryo-proper, while the PrE will form the endoderm of the visceral and parietal yolk sacs, and part of the embryonic gut tube (Nowotschin et al., 2019a). Though these lineages appear to be developmentally fixed, rare cells having committed to the EPI have been reported to switch to extra-embryonic endoderm, while the opposite has not been observed (Chan et al., 2019; Nowotschin et al., 2019b; Xenopoulos et al., 2015). Moreover, while PrE descendants contribute to the embryonic gut tube, they retain a partial transcriptional signature of their lineage of origin, suggesting a transcriptional, and perhaps epigenetic, memory (Kwon et al., 2008; Nowotschin et al., 2019b). These observations motivated us to seek a deeper understanding of how these two sister lineage identities are established and maintained.

A challenge in the study of early mammalian embryos is their relatively small size, limited cell number and availability. Alternative models, overcoming these limitations, are embryo-derived stem cells. Embryonic stem (ES) cells represent the pluripotent EPI (Evans and Kaufman, 1981; Martin, 1981), and extra-embryonic endoderm stem (XEN) cells represent the PrE (Kunath et al., 2005) (**Figure S1A**). Like the preimplantation EPI, ES cells express naïve pluripotency-associated markers, such as *Nanog*, *Oct4*, and *Sox2*, while XEN cells are defined by expression of PrE markers *Gata6*, *Gata4*, *Sox17* and *Pdgfra* (Garg et al., 2016; Watts et al., 2018). Additionally, upon reintroduction into the embryo, ES and XEN cells exclusively contribute cellular descendants to their lineage of origin, the EPI and PrE, respectively (Evans and Kaufman, 1981; Kunath et al., 2005; Martin, 1981).

Lineage conversion of *in vitro* stem cell models, along with cellular reprogramming, have been leveraged to examine the transcriptional and epigenetic mechanisms that control lineage identity, and determine the key milestones and bottlenecks in cell fate transition trajectories (Buganim and Jaenisch, 2012; Watts et al., 2018; Xu et al., 2015). Transcription factor (TF) modulation can be used to induce lineage conversions emphasizing the importance of TFs in lineage specification and maintenance. Accordingly, the endoderm-associated TFs *Gata4* and *Gata6* can convert ES cells to XEN cells (referred to as induced, or iXEN), which, along with observations made in mouse mutants (Bessonnard et al., 2014; Schrode et al., 2014), has established them as core members of the XEN/PrE gene regulatory network (Fujikura et al., 2002; Schröter et al., 2015; Shimosato et al., 2007; Wamaitha et al., 2015). However, TF-induced reprogramming in the reverse direction – the conversion of XEN cells to ES/iPS cells – has not been reported. Chemical reprogramming of somatic cells to induced pluripotent stem (iPS) cells has been reported to transition through a XEN-like state *en route* to successful reprogramming (Guan et al., 2022; Zhao et al., 2018, 2015), suggesting that XEN cells might represent a plastic state conferring the ability to acquire pluripotency. Alternatively, during TF-based reprogramming of somatic cells to iPS cells, emerging XEN-like cells have been described as a “dead-end” (Parenti et al., 2016; Schiebinger et al., 2019), suggestive of a refractory cell state. Therefore, it remains unclear whether XEN cells are amenable to attain a pluripotent state.

Here, we demonstrate that ectopic expression of TFs (OCT4, KLF4 and SOX2) can reprogram XEN cells to a stable pluripotent state. By contrast to the reciprocal GATA4-mediated ES-to-iXEN conversion, which occurs over 4 days and is >95% efficient, XEN-to-iPS conversion requires ∼3 weeks with an efficiency ∼0.2%, suggesting a differential developmental plasticity of these two sister lineages. Leveraging this *in vitro* reciprocal lineage conversion system, bookended by EPI (ES/iPS) and PrE (XEN/iXEN) states, we charted the sequence of events that drive cell state transitions, to identify facilitators or barriers of lineage switching. We applied single-cell transcriptomic analyses (scRNA-seq) to establish a high-resolution map of the EPI-PrE lineage conversions in both directions. These data revealed a linear, nearly deterministic, ES-to-iXEN conversion, and a heterogeneous and discontinuous XEN-to-iPS conversion with multiple intermediate terminal states along the trajectory. Dismantling of the XEN network by GATA4 knockout, but not GATA6, significantly increased the reprogramming efficiency of XEN-to-iPS. Although direct comparison between the XEN-to-iPS and ES-to-iXEN trajectories revealed some differences at key transition stages, comparison to *in vivo* scRNA-seq data from early mouse embryos, revealed that both *in vitro* trajectories mapped to *in vivo* trajectories during EPI and PrE lineage specification and maturation, but bypassed the ICM bipotent progenitor state. Bulk and single-cell assays for transposase-accessible chromatin by sequencing (scATAC-seq) revealed contrasting global levels and patterns of chromatin accessibility between ES/iPS and XEN/iXEN cells and suggested that extensive and late chromatin opening during XEN-to-iPS conversion represents a major roadblock. Altogether, these observations reveal that EPI and PrE have drastically different plasticities, and provide insights into the molecular basis of these differences.

## RESULTS

### OCT4, SOX2 and KLF4 can successfully reprogram XEN-to-iPS but in a slow and inefficient manner

To test the potential for XEN cells (**Figure S1A**), representing the PrE, to reprogram to iPS cells, we derived XEN cells from a transgenic doxycycline-inducible “reprogrammable” mouse strain used in previous studies for reprogramming of several somatic cell types, including mouse embryonic fibroblasts (MEFs) and various hematopoietic lineages (**Figure 1A**) (Bar-Nur et al., 2014; Stadtfeld et al., 2010). Doxycycline (‘dox’) induction with a constitutively expressed reverse tetracycline transactivator (Rosa26-M2rtTA) drives the expression of a polycistronic cassette consisting of *Oct4*, *Klf4*, *Sox2*, and an *mCherry* reporter (referred to as 3-factor, or ‘3F’). An additional *Pou5f1*-IRES-EGFP reporter allele enables detection of endogenous *Oct4* activation (*Oct4*-GFP), which is silent in XEN cells (Kunath et al., 2005; Lengner et al., 2007). Blastocysts harboring all three alleles were used to derive multiple XEN cell lines.

**Figure 1.**
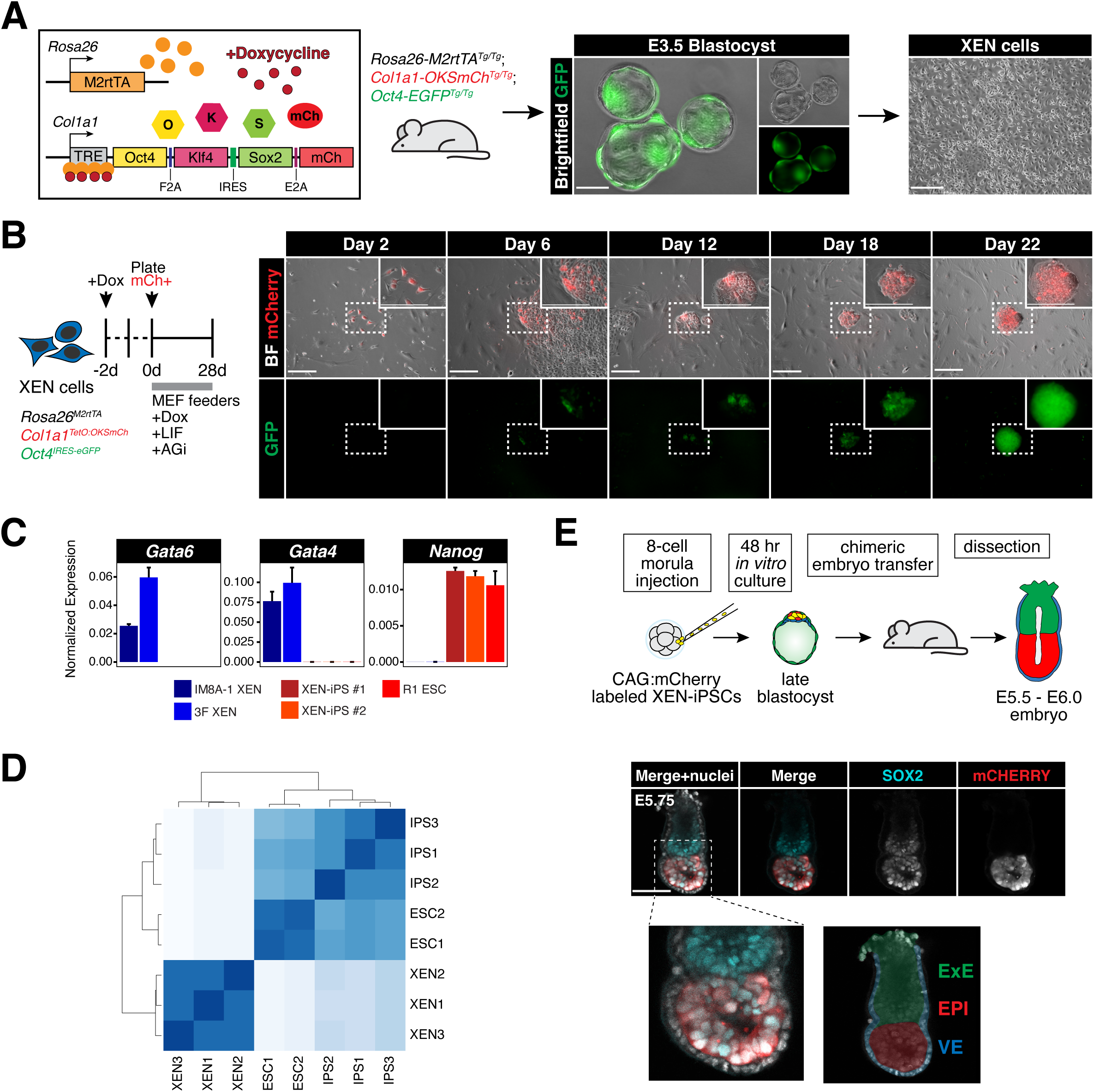
*Oct4*, *Sox2* and *Klf4* can successfully reprogram XEN cells but in a slow and inefficient manner. **(A)** Mice homozygous for 3 alleles (*R26:M2rtTA^Tg/Tg^*; *Col1a1:OKSmCh^Tg/Tg^*; *Oct4:EGFP^Tg/Tg^* referred to as ‘3F’) were intercrossed to collect blastocysts and derive XEN cells. These cells were then used to test their potential for reprogramming to iPS cells. **(B)** *(Left)* Experimental scheme representing the reprogramming conditions used for XEN cells (*see Methods for details*). *(Right)* Brightfield and fluorescent images of XEN cells during the reprogramming time course. Scale bars represent 250µm. **(C)** Gene expression data using RT-qPCR for several XEN and ES cell markers of wildtype XEN (IM8A-1), 3F XEN, two XEN-iPS lines (#1 and #2), and wildtype ES cells (R1). Individual bars show mean expression of three technical replicates normalized to mean expression of two reference genes: *Actb* and *Gapdh*; error bars represent standard deviation. **(D)** Unsupervised hierarchical clustering of bulk RNA-seq data from XEN-iPS cells, wild-type ES cells, and XEN cells. **(E)** *(Top)* Experimental scheme outlining the generation of XEN-iPS chimeric embryos and blastocyst transfers. XEN-iPS cells were labeled with mCherry before generating chimeras to track XEN-iPS contribution to the embryo at post-implantation stages. *(Bottom)* Maximum intensity projection of 5 optical sections from a confocal image of a chimeric embryo stained with the indicated markers. Nuclei were labeled using DAPI, mCherry was stained with an anti-RFP antibody. Scale bars represent 100µm.

Embryo-derived 3F XEN cells were treated with dox for 2 days prior to sorting of the mCherry+ fraction (∼15-30% of the total population) to ensure selection of cells expressing the reprogramming cassette. Sorted mCherry+ cells were plated for reprogramming in serum/LIF media with ascorbic acid and GSK3βi (‘AGi’), an enhanced reprogramming protocol which improves reprogramming efficiency in somatic cells, including conditions without exogenous *Myc* expression as in our experiments (Bar-Nur et al., 2014) (**Figure 1B**). Under these conditions, we were able to derive transgene-independent, iPS-like, *Oct4*-GFP+ colonies, although at very low frequency (<1%) and very slow kinetics (∼18-24 days). The derived XEN-iPS cells had silenced XEN/PrE markers including *Gata6*, *Gata4* and *Pdgfra*, and upregulated various pluripotency-associated markers, such as *Nanog*, *Esrrb*, *Fgf4*, *Zfp42* (*Rex1*), *Dppa3* (*Stella*) and *Utf1*, at levels similar to wild-type ES cells (R1), as detected by RT-qPCR and immunofluorescence analyses (**Figure 1C** and **S1B-C**). Hierarchical clustering and principal component analysis of bulk RNA-seq data clustered the XEN-derived iPS cells together with ES cell lines, and apart from the parental or published XEN lines (**Figure 1D** and **S1D**). To further assess developmental potential and lineage restriction of our XEN-iPS cells, we generated chimeric embryos by injecting them into 8-cell morula stage host embryos, prior to ICM lineage specification. A constitutively expressed CAG:mCherry construct was introduced into XEN-iPS cells to identify their descendants in post-implantation embryo chimeras (**Figure 1E**, top). Unlike their parental XEN cells (Kunath et al., 2005), XEN-iPS contributed exclusively to the epiblast compartment and were excluded from PrE-derived visceral endoderm and parietal endoderm tissues (**Figure 1E** and **S1E)**. These results demonstrate that TF-induced reprogramming of XEN cells can give rise to *bona fide* iPS cells, albeit in a slow and inefficient manner.

### Bidirectional reprogramming of XEN and ES cells reveals drastically different kinetics and efficiencies of conversion

To gain insights into the trajectory of XEN reprogramming toward iPS cells, and determine potential intermediate populations and rate limiting steps, we used flow cytometry to track the activation dynamics of two pluripotency-associated markers – the *Oct4*-GFP reporter present in our 3F XEN cells and the surface marker SSEA-1 (Cui et al., 2004; Lengner et al., 2007; Solter and Knowles, 1978) – as well as the silencing of the XEN marker PDGFRα (Artus et al., 2010; Plusa et al., 2008; Rugg-Gunn et al., 2012) (**Figure 2A** and **S2A**). We consistently observed asynchronous and independent activation of *Oct4*-GFP (O+) and SSEA-1 (S+) markers, with a small group of cells (∼0.5%) only expressing *Oct4*-GFP (O+) as early as day 4 of reprogramming, while another subset (∼0.1%) only expressed SSEA-1 at later stages (∼day 8). Cells co-expressing both pluripotency markers, SSEA-1 and *Oct4*-GFP (S+O+), were detected between days 14-16 of reprogramming. On the other hand, PDGFRα (P+) expression persisted in the majority of cells even after upregulation of *Oct4* and/or SSEA-1, and was downregulated only in the late stages of reprogramming, after ∼day 16. To determine whether acquisition or loss of any of these markers during reprogramming represented more advanced or delayed intermediates toward an iPS state, we sorted four different subpopulations at day 14 (S+O-P+, S+O+P+, S-O+P+ and S-O+P-) and re-plated them in reprogramming conditions (serum/LIF+AGi+dox) for 14 additional days prior to analysis (**Figure 2B**). Cells expressing SSEA-1 alone, or together with, *Oct4*-GFP (S+O- or S+O+) early on during reprogramming did not contribute significantly to the final S+O+P-iPS-like population (<5%), and often regressed to an S-state, suggesting that SSEA-1 is not a predictive marker of successful XEN reprogramming. However, cells expressing *Oct4*-GFP while silencing PDGFRα (S-O+P-) were the only intermediate population with significantly advanced reprogramming potential (>20%). These results document the slow and heterogeneous nature of XEN-to-iPS reprogramming, and suggest that silencing of the XEN program is a critical rate-limiting step.

**Figure 2.**
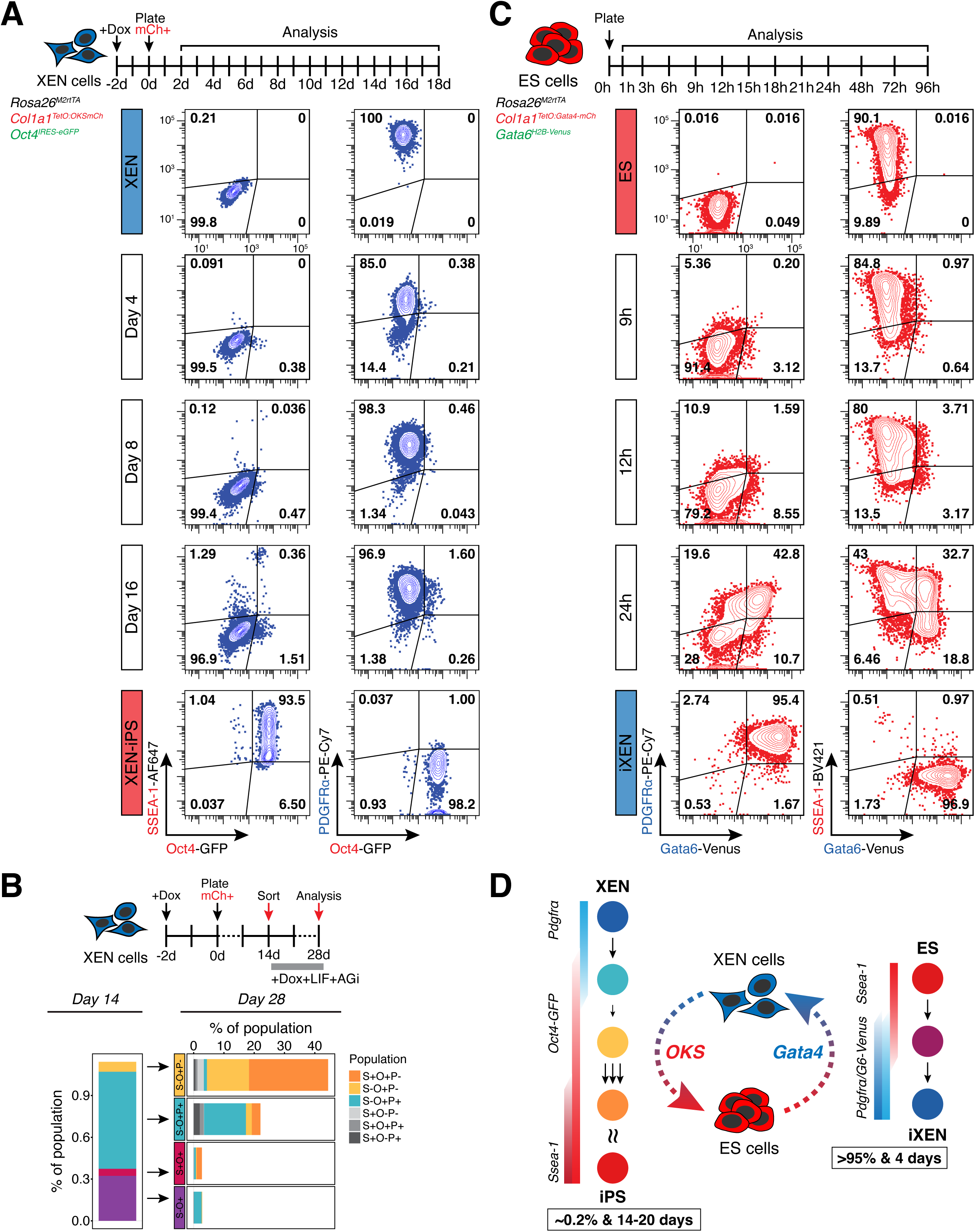
Reciprocal lineage conversions of XEN and ES cells have drastically different kinetics and efficiencies of conversion. **(A)** Time course tracking of XEN reprogramming using flow cytometry analysis of pluripotency-associated markers SSEA-1 and *Oct4*-GFP, and XEN-associated marker PDGFRα. *(Top)* Experimental scheme indicating analysis timepoints. *(Bottom)* Representative contour plots showing expression of SSEA-1 (AlexaFluor647-conjugated), *Oct4*-GFP and PDGFRα (PE-Cy7-conjugated) at days 2, 8 and 16 of reprogramming. Population percentage is indicated within each gate. **(B)** *(Top)* Experimental scheme describing timeline and methodology of tracking initial subpopulations that arise during XEN reprogramming. Four major subpopulations were sorted at day 14 of reprogramming. Following 14 additional days of reprogramming (day 28), populations arising from each sorted subpopulation were determined using flow cytometry. *(Bottom, left)* Stacked bar charts depicting the mean proportion of the entire population represented by each subpopulation at day 14, and at day 28 *(bottom, right)*. S-O-P+ subpopulation constitutes the remainder of the population (not shown) to amount to 100% and is indicative of XEN cells that do not change their expression status of either marker assayed in this experiment compared to the starting state. N = 6 (2 independent experiments consisting of 3 replicates each). **(C)** Time course tracking of ES-to-iXEN conversion using flow cytometry analysis of pluripotency-associated markers SSEA-1, and XEN-associated markers PDGFRα and *Gata6*-Venus. *(Top)* Experimental scheme indicating analysis timepoints. *(Bottom)* Representative contour plots show expression of PDGFRα (PE-Cy7-conjugated), *Gata6*-Venus and SSEA-1 (BV421-conjugated) at indicated timepoints. Population percentage is indicated within each gate. **(D)** Summarized schematic of XEN-to-iPS and ES-to-iXEN lineage conversions. *(Left)* Hypothesized reprogramming route taken by XEN cells. Sizes and number of arrows reflect the likelihood of cells progressing from one state to the next (smaller/single arrows = few cells progress; larger/more arrows = many cells progress). Based on the population tracking data in panels A-B, XEN cells initially express *Oct4*-GFP before downregulating expression of PDGFRα. This downregulation step appears to be a bottleneck in the reprogramming process since very few *Oct4*-GFP+ cells progress to this state. Following downregulation of PDGFRα, a large proportion of cells will upregulate SSEA-1 expression. *(Right)* Hypothesized route of lineage conversion taken by ES cells. Initial upregulation of both endoderm markers – PDGFRα and *Gata6*-Venus is followed by downregulation of SSEA-1. No obvious bottlenecks are detected during ES-to-iXEN conversion.

Although the exact reprogramming efficiency and kinetics varied with different 3F XEN lines, they remained consistently low (∼0.2%) and slow (∼20 days) across all lines tested (**Figure S2B**). In contrast, the opposite cell fate transition from ES to iXEN has been reported to be very efficient (McDonald et al., 2014; Schröter et al., 2015; Wamaitha et al., 2015). To directly compare reciprocal interconversion efficiencies and trajectories, we used an ES cell line engineered to express *Gata4*-mCherry fusion protein in a dox-inducible manner (Schröter et al., 2015). This system enables a quick and efficient conversion to a XEN-like state (induced XEN or iXEN) over the course of 4 days, with >95% of cells silencing pluripotency markers (e.g. SSEA-1), and activating XEN markers, including PDGFRα and endogenous *Gata6* linked to a H2B-Venus reporter (*Gata6*-Venus) (Freyer et al., 2015; Schröter et al., 2015) (**Figure 2C-D** and **S2A**). Although activation of XEN-associated PDGFRα and *Gata6*-Venus occurs as early as ∼9-12h, most cells continue to co-express the pluripotency-associated marker SSEA-1, which is subsequentially silenced.

Together, these results document a prominent difference in the efficiency and kinetics of ES and XEN cell interconversions. Transitioning from a XEN-to-iPS state is slow and involves a heterogeneous trajectory, in contrast to the opposite ES-to-iXEN conversion. This argues that XEN cells are “locked” into a less plastic state with a stable transcriptional program that is resistant to reprogramming and silencing, as documented by the persistent expression of XEN markers until the very late stages of conversion. In further agreement, nascent iXEN cells derived from iPS through transient *Gata4* expression show a diminished tendency to re-acquire an iPS state by dox-inducible OKS expression – with reprogramming efficiencies similar to embryo-derived XEN cells (**Figure S2C**). Notably, despite their distinct kinetics and efficiencies, conversions in both directions (ES-to-iXEN and XEN-to-iPS) appear to transition through a state where both ES and XEN markers are co-expressed, suggesting an, at least partially, overlapping trajectory.

### Single-cell transcriptomics reveals a discontinuous XEN-to-iPS reprogramming trajectory with multiple end-states in contrast with the linear ES-to-iXEN conversion

Noting the heterogeneities observed, we sought to map the trajectories of these bidirectional cell fate transitions at a single-cell level. We profiled cell states using scRNA-seq at five distinct timepoints during each conversion – embryo-derived XEN or ES cells and fully reprogrammed iPS or iXEN cells –representing starting and end states, as well as cells from early, middle and late stages of reprogramming in each direction (**Figure 3A**). Samples were collected in technical duplicates, with approximately 8,000 cells sampled from each replicate and timepoint (i.e., ∼16,000 cells total per timepoint; ∼160,000 cells total across all timepoints and both trajectories). Since XEN reprogramming to iPS cells is an inefficient process, to avoid potential underrepresentation of cells undergoing successful reprogramming, which constitute only a small fraction of the bulk population, we sorted four intermediate subpopulations based on SSEA1 and *Oct4*-GFP expression (see **Figure 2B** and **S2A**) – S-G-, S+G-, S+G+ and S-G+ – and reconstituted them in roughly equal proportions prior to droplet encapsulation (**Figure S3A**). We followed this approach for all three intermediate timepoints for the XEN-to-iPS trajectory, representing day 7, day 14 and day 28 of reprogramming. A similar strategy was used for the early (9h) timepoint for the ES-to-iXEN trajectory (**Figure S3A**).

**Figure 3.**
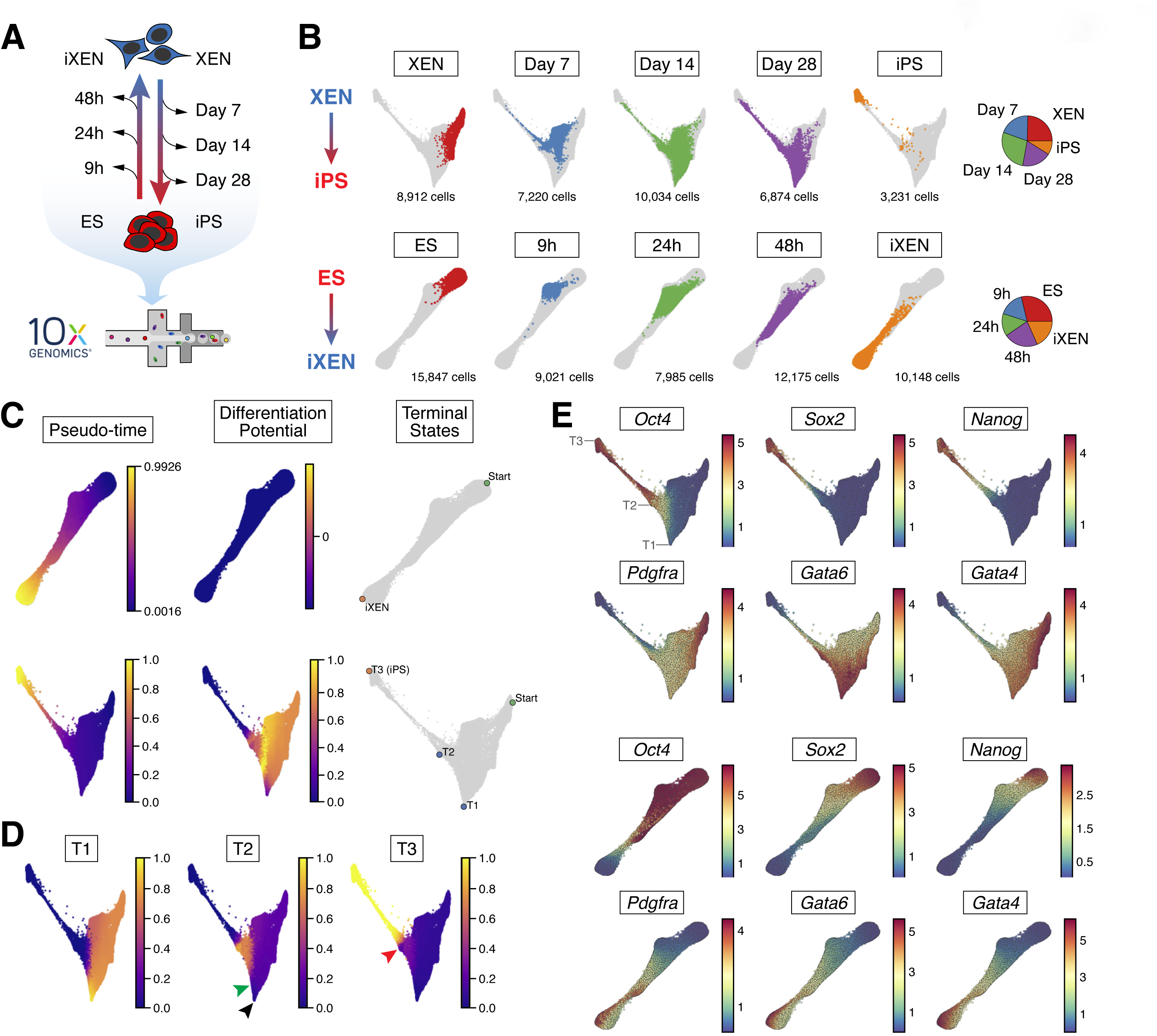
scRNA-seq analyses of XEN-to-iPS and ES-to-iXEN conversions. **(A)** Experimental scheme outlining the cells and lineage conversion timepoints assayed by scRNA-seq. **(B)** Force-directed layouts showing pooled timepoints of XEN-to-iPS *(top)* and ES-to-iXEN *(bottom)* conversion in a single trajectory. Individual plots highlight the distribution of single timepoints across the trajectory. *(Right)* Pie chart representations of the proportion of cells representing each collection timepoint relative to the entire cohort per conversion trajectory. **(C)** Palantir determined pseudotime ordering, terminal states, and differentiation potential of ES-to-iXEN *(top)* and XEN-to-iPS *(bottom)* trajectories. **(D)** Branch probabilities of terminal states determined by Palantir in the XEN-to-iPS trajectory. Black arrowhead indicates cells at T1 with low probability of differentiating to T2. Green arrowhead indicates where T2 probability increases, coinciding with *Oct4* expression. Red arrowhead indicates cells at T2, where they have a non-zero probability of acquiring the T3 state. **(E)** Gene expression patterns of XEN and pluripotency-associated markers. Each cell is colored on the basis of its MAGIC imputed expression level for the indicated gene. Locations of T1, T2 and T3 terminal states are indicated for the XEN-to-iPS trajectory.

To gain insights into when and how lineage conversions occur, we combined data from all replicates and timepoints separately for each reprogramming trajectory and projected them on a Force Directed Layout (FDL) (**Figure 3B**). We then applied Palantir (Setty et al., 2019), to automatically identify the terminal states of the system, and the branch probabilities of each cell reaching each of the identified states. The differentiation potential (entropy) of these branch probabilities represents the uncertainty of future cell fate (**Figure 3C** and **3D**). Consistent with the rapid and efficient dynamics of ES-to-iXEN conversion, we identified a linear trajectory and a singular terminal state that cells eventually reach in this trajectory (**Figure 3B** and **3C**). By contrast, during XEN reprogramming we detected three distinct terminal states, ‘T1’, ‘T2’ and ‘T3’, highlighting the inefficient and more heterogeneous nature of XEN-to-iPS conversion. The most prominent T1 state represented a XEN-like state with high expression of XEN genes and absence of pluripotency gene expression, but also with upregulation of AP-1 family members (such as *Jun, Fos, Atf3*), which have been previously shown to inhibit somatic cell reprogramming in different contexts (Chronis et al., 2017; Liu et al., 2015; Markov et al., 2021)(**Figure 3E** and **S3B-C**; **Table S1**). On the other hand, T3 reflected the iPS/EPI state characterized by expression of pluripotency-associated genes and silencing of the XEN/PrE-program. Finally, the T2 state was defined by expression of multiple XEN genes along with some pluripotency-associated markers, such as *Oct4* (endogenous), and *Rex1* (Nichols et al., 1998; Rogers et al., 1991; Schöler et al., 1989), in agreement with our flow cytometry data showing transient co-expression of the EPI and PrE markers during XEN reprogramming.

Palantir inferred that cells at the start of the trajectory had high probability of reaching T1 (**Figure 3D**). However, cells at T1 showed negligible probability of acquiring the T2 or T3 states, indicating that T1 serves as a ‘sink/dead-end’ during XEN-to-iPS reprogramming (**Figure 3D**, black arrowhead). These results suggest that the inefficiency of reprogramming is in part due to cells being diverted toward a stable and refractory T1 state. However, cells at T2, showed a low but non-zero probability of reaching the T3 state (red arrowhead), indicating that T2 is also a bottleneck toward successful reprogramming, but represents an intermediate more plastic state compared to T1. Once cells passed the T2 state, the probability of reaching T3, the end-point reprogrammed state, sharply increased. This phase of the trajectory was characterized by a downregulation of XEN-associated genes accompanied by the activation of additional pluripotency-related genes (**Figure 3E** and **S3C**). Notably, *Oct4* (as well as other pluripotency regulators, e.g., *Rex1* and *Klf9*) and *Dnmt3l* were upregulated prior to the T2 state, and their activation coincided with an increase in the probability of cells reaching T2 (**Figure 3D** and **3E**, green arrowhead), suggesting that this is an important but not sufficient step for XEN-to-iPS reprogramming in agreement with our sorting analyses (see **Figure 2B**).

In sum, our scRNA-seq data support XEN-to-iPS reprogramming as a heterogeneous process with multiple terminal states representing major roadblocks that must be bypassed to silence the PrE program and establish the EPI state.

### XEN and ES reprogramming approximate in vivo cell states, but not ICM progenitor cells

The drastically different efficiencies of the two lineage interconversions might suggest that they transition through unique intermediate states. To examine this possibility, we used Harmony (Nowotschin et al., 2019b) to combine all transcriptomes from XEN-to-iPS and ES-to-iXEN trajectories to derive a common reduced dimensional space (**Figure 4A-C)**. Merging of all datasets indicated that while the starting and end states of the two trajectories (ES/iPS and XEN/iXEN) are similar, the trajectories show differences during the transition stages of lineage conversion. To quantify this difference, we first utilized the individual trajectories to bin cells along the XEN-to-iPS and ES-to-iXEN conversions, respectively (**Figure S4A-B**). We then computed the average phenotypic distance between each pair of bins and visualized the pairwise distance matrix as a heatmap (**Figure S5A**), which illustrated the similarity at the terminal points of the transition and dissimilarity at intermediate stages. To further understand this dissimilarity, we computed the differentially expressed genes at intermediate states during each conversion. For this, we first aligned the two trajectories using a different Harmony algorithm (Korsunsky et al., 2019), and clustered the cells at the intermediate stages, followed by differential expression analysis using MAST (Finak et al., 2015) (**Figure S5B-C**; **Table S2**). Our analysis revealed that genes upregulated in the ES-to-iXEN trajectory included chromatin modifiers, such as *Jarid2*, *Hmga2* and *Arid1a*, while genes expressed in the XEN-to-iPS showed enrichment of mitochondrial and cellular metabolism-related genes. These findings indicated that reprogramming between EPI and PrE lineages involves transitioning through divergent cellular states, likely driven by unique epigenetic and metabolic regulators associated with the starting XEN and ES states (Gatie and Kelly, 2018; Gatie et al., 2022; Mulvey et al., 2015; Rugg-Gunn et al., 2010; Senner et al., 2012). Additionally, transition cell states during XEN-to-iPS reprogramming upregulated genes encoding several members of the AP-1 complex, such as *Atf3*, *Fos*, *Jun*, *Junb*, likely reflecting the “refractory” T1 state that cells acquired during this conversion. We confirmed these findings with an alternative batch alignment method using Harmony (Nowotschin et al., 2019b) and Spectral Clustering for grouping cells (see Methods for details).

**Figure 4.**
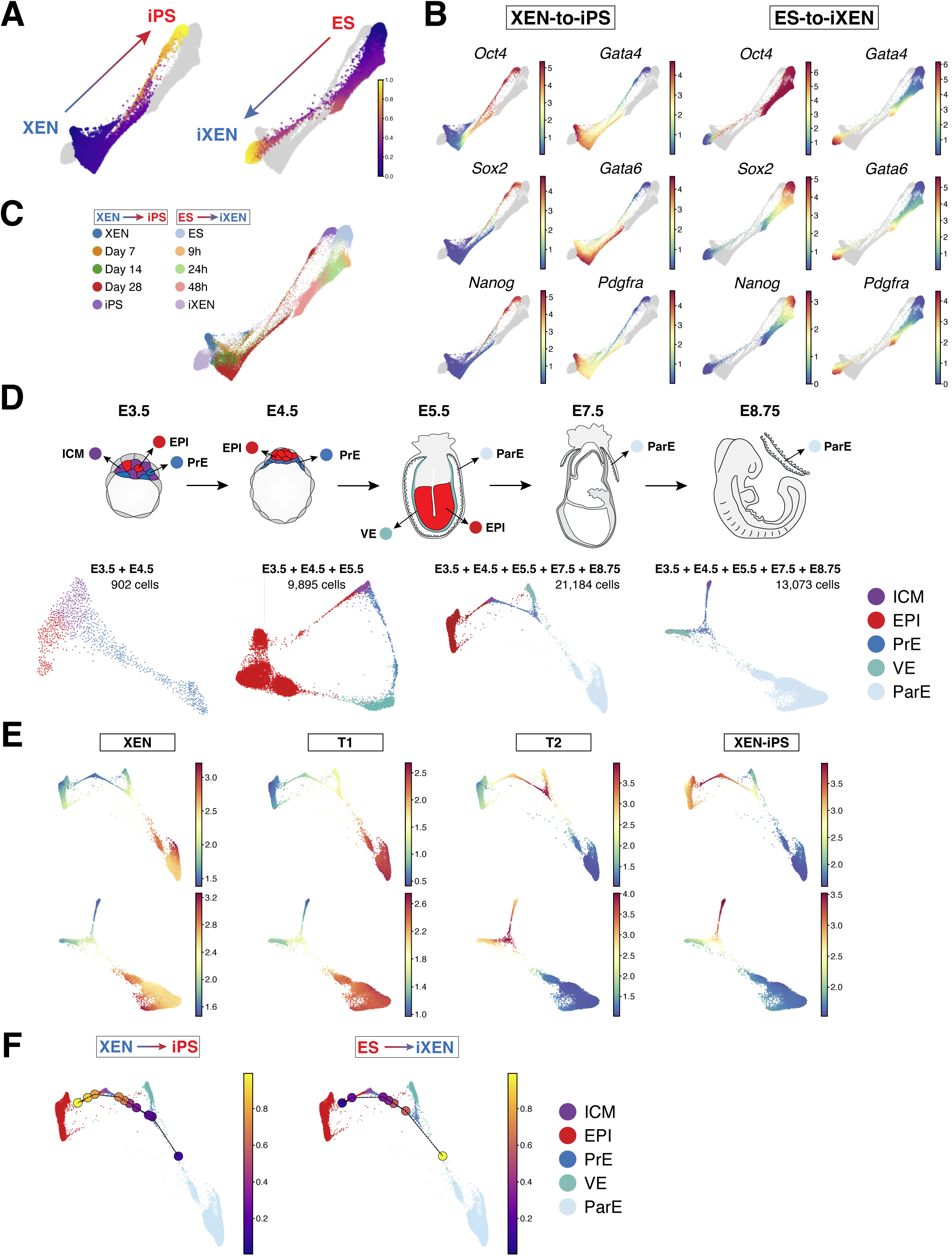
XEN-to-iPS and ES-to-iXEN conversion trajectories approximate *in vivo* cell states. **(A)** Force-directed layout of combined XEN-to-iPS and ES-to-iXEN trajectories based on Harmony (Nowotschin et al., 2019b) integration. XEN-to-iPS *(left)* or ES-to-iXEN *(right)* are highlighted. Cells are colored by Palantir pseudotime, computed separately as in Figure 3. **(B)** Gene expression patterns of various pluripotency and XEN-associated markers displayed in the combined trajectory. Each cell is colored on the basis of its MAGIC imputed expression level in the individual XEN-to-iPS or ES-to-iXEN trajectories, as labeled, for the indicated gene. **(C)** Force-directed layout of combined XEN-to-iPS and ES-to-iXEN trajectories (as in panel A) with individual time points from each trajectory colored as indicated. **(D)** *(Top)* Schematic illustrating *in vivo* embryo stages and tissues (labeled) profiled by scRNA-seq. *(Bottom)* Force-directed layouts of combined *in vivo* stages and lineages as labeled above and color-coded as indicated by cell type. **(E)** Gene expression signatures of XEN, XEN-iPS and T1 and T2 terminals states mapped onto force-directed layouts of combined *in vivo* states including the EPI lineage *(top)* or excluding the EPI *(bottom)*. **(F)** Visualization of *in vitro* XEN-to-iPS bins *(left)* or ES-to-iXEN bins *(right)* mapped onto the combined trajectory of *in vivo* bins. Individual dots represent single *in vitro* bins and are color coded according to pseudotime.

We next sought to determine the degree to which the *in vitro* reprogramming trajectories resembled cell states *in vivo* during the emergence and differentiation of pluripotent EPI and PrE lineages in the embryo. For this, we took advantage of our published scRNA-seq datasets from early preimplantation (E3.5 and E4.5) mouse embryos (Nowotschin et al., 2019b) to compile a reference subset of *in vivo* cells comprising uncommitted ICM progenitors, their derivative EPI and PrE cell lineages (**Figure 4D** and **S5D**), as well as the subsequent EPI and visceral endoderm (VE) present at the early post-implantation stage (E5.5). Since XEN cells have been suggested to resemble the parietal endoderm (ParE) branch of the PrE lineage due to their morphology, marker expression and lineage contribution in embryo chimeras (Artus et al., 2012; Brown et al., 2010; Kruithof-de Julio et al., 2011; Kunath et al., 2005), we supplemented our published embryo-derived atlas with newly generated scRNA-seq data of ParE cells collected from E7.5 and E8.5 embryo parietal yolk sacs.

We first used Harmony (Nowotschin et al., 2019b) to aggregate the *in vivo* timepoints, then used the combined *in vivo* trajectories, with or without the EPI lineage (i.e., highlighting the endoderm lineage), to identify the nearest *in vivo* states resembling the starting and terminal states (T1, T2 and T3/iPS) of the XEN-to-iPS conversion (**Figure 4D** and **S5D**). We visualized the average expression of genes that were significantly differentially expressed in each of the terminal states (**Figure 4E**). The starting XEN state mapped predominantly to the ParE in the combined *in vivo* trajectory, providing an unbiased transcriptional basis for its ParE-like character, and the observed preferential ParE contribution of XEN cells when reintroduced into embryos (Kunath et al., 2005). The terminal T1 state also mapped to the ParE, supporting the notion that T1 represents a stable state refractory to successful reprogramming (see **Figure 2E**). In contrast, T2 cells showed closer proximity to the PrE indicative of their progression away from the starting ParE state during reprogramming. Finally, T3, or XEN-iPS, cells mapped to the EPI as expected of pluripotent stem cells.

We next mapped the entire *in vitro* XEN-to-iPS and ES-to-iXEN trajectories on the *in vivo* datasets to determine their progression along *in vivo* developmental states during lineage conversion. We used Harmony (Nowotschin et al., 2019b) to combine all *in vivo* and *in vitro* cell transcriptomes to derive a common augmented representation. The *in vivo* and *in vitro* trajectories were separately divided into equal-sized pseudotime bins and the closest *in vivo* bin for each *in vitro* bin was identified in the augmented representation (**Figure S4** and **S5E-F**). In agreement with the previous analysis, we noted that the similarity to ParE persists from the starting XEN to the T1 state. As cells progressed to the T2 state during reprogramming they approximated the *in vivo* PrE, particularly at E4.5, progressing to E3.5 PrE, before moving to the EPI at E3.5 and, eventually, E4.5 EPI (**Figure 4F**). Cells undergoing ES-to-iXEN conversion followed a similar but opposite path along the *in vivo* trajectory, though not identical, reflecting their differences noted previously, but suggesting that both conversions bore resemblance to cell states present *in vivo* (**Figure 4F** and **S5F**). Notably, neither the XEN-to-iPS nor ES-to-iXEN trajectories transitioned through a state that resembled an uncommitted ICM progenitor. These observations were consistent irrespective of the size and number of bins used, and this combined with the high resolution of our scRNA-seq data revealed that EPI and PrE interconversions *in vitro* do not transition through an ICM-like state. Moreover, our observed toggling of cells between the EPI and PrE branches lends support to a model of bistability of the ICM lineages over tristability of EPI, PrE and ICM states, as has been suggested in studies modeling ICM development (Bessonnard et al., 2014; De Mot et al., 2016; Saiz et al., 2020; Schröter et al., 2015).

### Silencing of the XEN program is a bottleneck for successful XEN-to-iPS reprogramming

Having noted the discontinuous and inefficient nature of reprogramming XEN-to-iPS states, we wanted to further explore the transcriptional determinants and roadblocks of this lineage conversion. We first tracked gene expression changes along pseudotime to identify genes that were down/upregulated early versus late in the process (**Figure 5A** and **S6A**; **Table S3**). Individual genes were clustered based on similarity of their expression trends along pseudotime (gene clusters with fewer than 20 genes are not shown). We noticed that pluripotency-associated genes followed distinct trends of early, gradual or late upregulation along pseudotime. In agreement with our flow cytometry data, *Oct4* was among the early activated genes, which also included *Rex1*, *Mybl2*, *Crxos*, *Egr1* and *Fgf4* (Cluster 0), while other key pluripotency regulators such as *Nanog* and *Esrrb* (Clusters 8 and 5, respectively) were only upregulated at the final stages, suggesting the presence of barriers for their activation. In contrast with the asynchronous activation of pluripotency-associated genes, the endoderm-related transcriptional program was silenced in a slow but coordinated manner with an initial downregulation around the T2 stage but complete silencing only achieved during the final stages of reprogramming. We therefore hypothesized that dismantling the XEN network was a critical milestone for successful XEN-to-iPS reprogramming. In agreement, flow sorted cells having lost PDGFRα expression (S-O+P-) at day 14 were able to generate stable, transgene-independent iPS-like cells, when plated in serum/LIF medium in the absence of dox and AGi, while cells with persistent PDGFRα expression (S-O+P+) failed to do so (**Figure S6B**).

**Figure 5.**
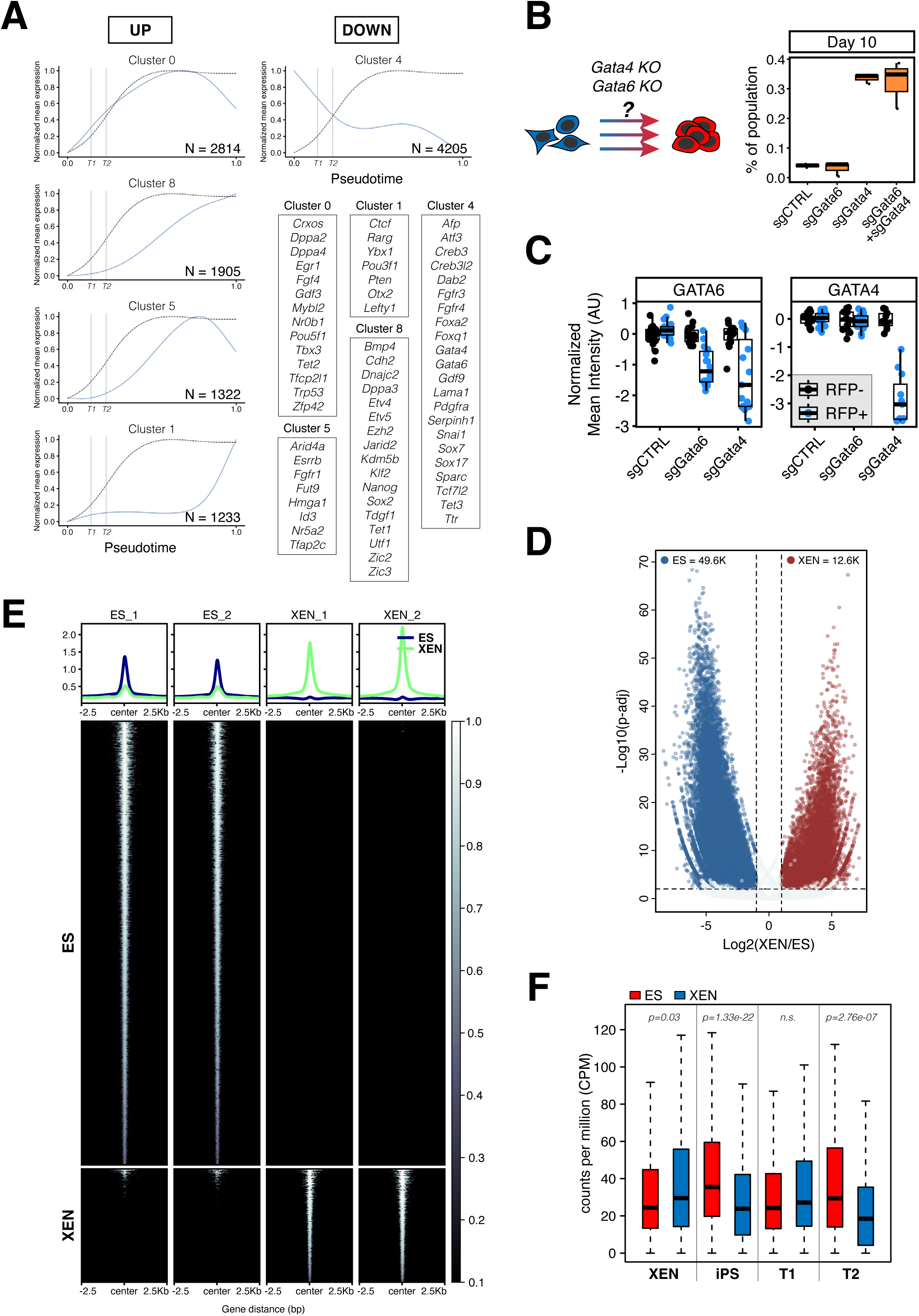
XEN transcriptional network serves as a roadblock to successful XEN-to-iPS reprogramming. **(A)** Gene expression waves over pseudotime of XEN-to-iPS reprogramming. Plots show mean expression trend of all genes within each cluster. Dotted curve represents the probability of acquiring the T3/iPS state. Vertical dotted lines indicate T1 and T2 terminals states along the pseudotime axis. Representative genes for each cluster are highlighted in boxes. **(B)** Reprogramming efficiency following *Gata4* and/or *Gata6* perturbation. Reprogrammable XEN cells were transfected with plasmids constitutively expressing Cas9 and sgRNAs targeted to *Gata4* and/or *Gata6*, or control sgRNA. Cells were plated in reprogramming conditions for 10 days and resulting percentage of iPS-like cells was determined using flow cytometry. *(Right)* Box plots showing the proportion of the entire population that were S+O+P-at day 10 of reprogramming. Middle line marks the median; lower and upper hinges correspond to the first and third quartiles, respectively. Whiskers extend to 1.5*interquartile range (IQR) from the hinge. Outliers are represented by open circles. N = 3. **(C)** Box plots showing relative reduction in anti-GATA4 or anti-GATA6 fluorescence immunostaining in XEN cells transfected with Cas9/sgRNA expression vectors targeting *Gata4* or *Gata6*, or non-target control. Individual points represent relative fluorescence intensity in individual cells represented as arbitrary units and normalized to untransfected (RFP-) cells within the same well (see images in Figure S6C). **(D)** Volcano plot of significantly differential accessible ATAC-seq peaks in XEN versus ES cells. **(E)** Tornado plots of ATAC-seq signal in XEN and ES cells. The ATAC-seq signals are shown for 2.5kb up- and downstream of peak centers. **(F)** Box plots showing relative chromatin accessibility in XEN and ES cells of genes enriched in XEN, XEN-iPS, T1 and T2 terminal states.

To directly test the inhibitory effect of the XEN program on XEN-to-iPS conversion, we knocked out *Gata4* and/or *Gata6,* two master regulators of the PrE state, and determined the impact on XEN-to-iPS reprogramming. Both factors are potent inducers of the XEN state in ES cells; GATA6 is required for PrE specification *in vivo* and downstream activation of *Gata4* expression, while GATA4 is required for PrE lineage differentiation (Bessonnard et al., 2014; Fujikura et al., 2002; Schrode et al., 2014; Schröter et al., 2015; Shimosato et al., 2007; Wamaitha et al., 2015). Using CRISPR-Cas9 technology, we knocked out *Gata4* or *Gata6* individually and together (**Figure S6C**) in XEN cells prior to reprogramming and assessed the relative efficiency of reprogramming by measuring the proportion of S+O+P-cells arising in culture at day 10 of XEN-to-iPS reprogramming (**Figure 5B**). While *Gata4* deletion improved reprogramming efficiency by ∼7-fold, *Gata6* deletion did not show a significant increase in iPS-forming potential. Additionally, deletion of both factors did not have a synergistic effect over deletion of *Gata4* alone. Finally, *Gata4* KO induced a notable reduction in *Gata6* expression, but not vice versa (**Figure 5C** and **S6C**). Together, these data demonstrate that overwriting the PrE transcriptional program is a major bottleneck for successful XEN-to-iPS reprogramming and identify GATA4, but not GATA6, as a dominant factor involved in maintenance of the PrE program in XEN cells.

### Remodeling to an EPI-like chromatin state constitutes another roadblock during XEN-to-iPS reprogramming

One explanation for the drastically different efficiencies of XEN and ES interconversions could be the differential epigenetic plasticity of the two lineages. ATAC-seq analysis in bulk ES and XEN cells, revealed a significantly higher proportion of loci exhibiting open chromatin in ES cells as compared to XEN cells (49625 versus 12670, respectively), suggesting that the ES cell genome is generally more accessible than the XEN cell genome (**Figure 5D-E**). Additionally, we noted that several XEN-associated loci (XEN and T1 signature genes based on the scRNA-seq analysis) showed similar levels of accessibility in XEN and ES cells, while iPS/ES-associated loci (iPS signature genes) exhibited significantly lower accessibility in XEN cells (**Figure 5F** and **S6D**). Notably, genes associated with the T2 state also showed reduced accessibility in XEN cells, compared with ES/iPS cells. Based on these observations, we wondered whether chromatin remodeling towards a more open and plastic EPI state could represent another roadblock impacting XEN reprogramming.

To address this question, we performed single-cell ATAC-seq (scATAC-seq) to detect and quantify dynamic accessibility changes during reprogramming. We sampled ∼100,000 nuclei from the same timepoints during lineage conversion in either direction (XEN-to-iPS and ES-to-iXEN) as in our scRNA-seq dataset, and generated ‘metacells’ using the SEACells algorithm (Persad et al., 2022). Individual metacells represent small groups of single cells (110 on average) with similar accessibility states (see Methods for details). We focused our analysis of the XEN-to-iPS conversion on loci with highly variable accessibility among metacells (N=3584; **Table S4**). A large fraction of these loci (40.49%) represented ES-specific peaks (compared to only 4.63% overlapping with XEN-specific peaks) as detected by bulk ATAC-seq (presented in **Figure 5E**) and overlapped predominantly with putative enhancers of ES cells compared with XEN, as detected by bulk H3K27ac ChIP-seq (31.36% versus 4.49%, respectively) (**Figure 6A**). These results indicate that XEN-to-iPS reprogramming is accompanied by extensive chromatin opening around EPI-related regulatory regions.

**Figure 6.**
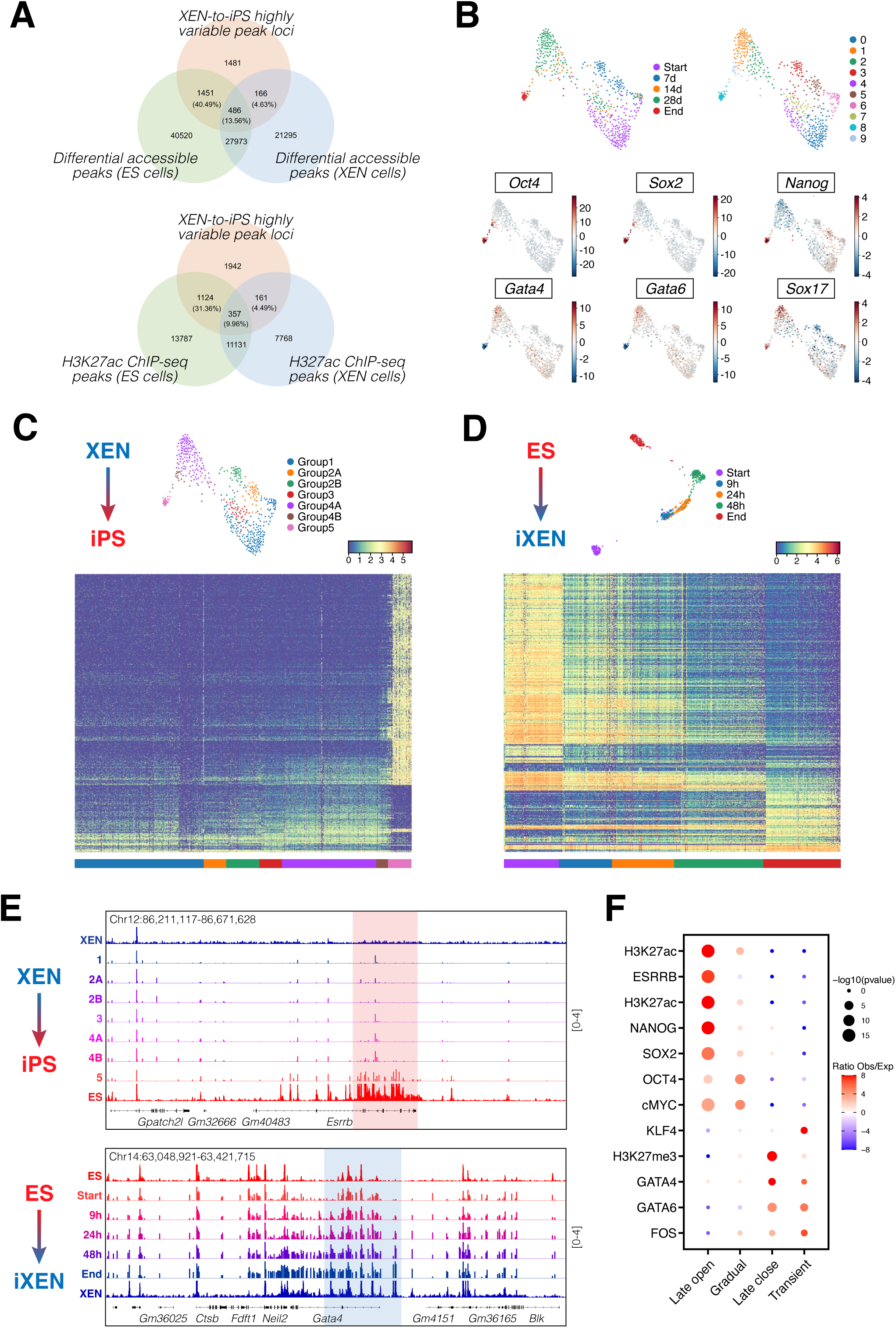
Establishing an EPI-like chromatin state underlies the inefficient conversion of XEN to iPS cells. **(A)** Venn diagrams depicting the number and percentages of highly variable accessible peak loci identified from scATAC-seq data for the XEN-to-iPS conversion trajectory that overlap with differential accessible peak loci identified from bulk ATAC-seq data for XEN and ES cells *(top)*, or with H3K27ac ChIP-seq peak loci in XEN and ES cells *(bottom)*. **(B)** Force-directed layouts showing the combined scATAC-seq dataset for XEN-to-iPS conversion and highlighting the individual timepoints *(top left)*, metacell clusters *(top right)*, or ChromVAR scores for XEN- and ES-specific TFs *(bottom)*. **(C)** *(Top)* Force-directed layout showing the scATAC-seq dataset for XEN-to-iPS conversion and highlighting the groups of metacells identified based on similar accessibility profiles along pseudotime. *(Bottom)* Heat map view of the relative chromatin accessibility changes over pseudotime of XEN-to-iPS conversion for a subset of selected peak loci, going from group 1 to group 5 of the metacells. Each row corresponds to a peak locus and each column corresponds to a metacell. The peak loci shown are the differential accessible peak loci identified in the XEN-to-iPS conversion based on differential accessibility analysis of peak loci between group 4B and group 5 of the metacells. **(D)** *(Top)* Force-directed layout showing the scATAC-seq dataset for ES-to-iXEN conversion and highlighting the individual timepoints. *(Bottom)* Heatmap view of the relative chromatin accessibility changes of ES-to-iXEN conversion based on the highly variable peak loci that overlap between the ES-to-iXEN and XEN-to-iPS conversions. **(E)** Example IGV (Integrative Genomics Viewer) tracks showing accessibility peaks of pseudo-bulk scATAC-seq data of XEN-to-iPS conversion *(top)* or ES-to-iXEN *(bottom)*. Highlighted are relative accessibility in metacell groups (top panel; XEN-to-iPS) or individual timepoints (bottom panel; ES-to-iXEN) at example genomic loci showing late opening in XEN-to-iPS reprogramming, versus gradual opening in ES-to-iXEN conversion. Signal values are indicated to the right. **(F)** LOLA enrichment analysis of late opening, gradual opening, late closing or transient opening peaks during XEN-to-iPS conversion.

ChromVAR analysis depicting accessibility changes around the motifs of key EPI or XEN-related regulators, suggested that key cell identity-related changes occurred mostly during the final stages of reprogramming. Notably, motifs for OCT4 and SOX2 – despite their continuous exogenous expression – displayed high accessibility only at the late stages of XEN-to-iPS reprogramming (**Figure 6B**). For a more unbiased analysis of the kinetics and potential drivers of accessibility changes, we performed PhenoGraph (Levine et al., 2015) clustering of scATAC-seq metacells undergoing XEN-to-iPS conversion, which assigned 7 major groups (1, 2A, 2B, 3, 4A, 4B and 5) based on relative accessibility along lineage conversion (**Figure 6C**). A heatmap of the most variable peaks (N=3584; **Table S4**) across the 7 groups revealed an extensive and slow chromatin opening and rather restrictive closing during XEN-to-iPS conversion, in agreement with the higher chromatin accessibility of the EPI compared to the PrE state. On the other hand, tracking the most variable peaks during the opposite ES-to-iXEN transition (N=1955 that overlap with the highly variable peaks of the XEN-to-iPS trajectory; **Table S5**) revealed more rapid and progressive changes (**Figure 6D-E** and **S7**), consistent with the more efficient nature of this conversion. Enrichment analyses using published ChIP-seq datasets (Sheffield and Bock, 2016) showed a significant association for known regulators of EPI (e.g. NANOG, OCT4, SOX2, ESRRB) or PrE fate (e.g. GATA4, GATA6) among the opened or closed regions, respectively (**Figure 6F; Table S6**). However, we also detected a subset of regions that opened more gradually in the XEN-to-iPS trajectory, as well as a group of transiently accessible regions that only lost accessibility once pluripotency was established (**Figure 6C**). Gradually opening peaks were enriched for binding of KLF4 and MYC, while transiently open peaks were enriched for GATA factors (PrE program) and FOS (AP-1 complex) binding, and overlapped with sites that are silenced by H3K27me3 histone marks in ES cells. This could suggest that the persistent expression of GATA factors might derail reprogramming by opening up new sites toward alternative fates or dead-end states. Together, these data reveal that many critical EPI and PrE regulatory elements are refractory to chromatin remodeling during the XEN-to-iPS conversion, likely contributing to this being a slow and inefficient process.

In conclusion, by applying single-cell analyses on a bidirectional reprogramming system, we were able to demonstrate that ES and XEN cells, representing the sister EPI and PrE lineages of the embryo, have drastically different interconversion efficiencies, suggesting differential plasticities. The inefficient and slow XEN-to-iPS conversion is highlighted by the dominant PrE transcriptional program, which is partly governed by GATA4, and is challenging to overwrite in the presence of potent EPI-associated factors. Moreover, the extensive chromatin remodeling required by XEN cells to acquire the widely accessible chromatin state of ES cells likely imposes an additional roadblock to successful reprogramming. Finally, by mapping the ES-to-iXEN and XEN-to-iPS *in vitro* trajectories on the scRNA-seq trajectories from developing embryos, we showed that cells progressing in either direction tracked along and toggled between EPI and PrE *in vivo* state trajectories without transitioning through an ICM progenitor state, underscoring a bistable system.

## DISCUSSION

The EPI and PrE lineages arise from the common ICM progenitor, and thus have a common lineage history and unique lineage relationship during mammalian development, the molecular basis of which remains poorly understood. We therefore sought to probe the relative plasticity of these two sister lineages and determine whether PrE cells can acquire a pluripotent state by manipulating the transcription factor network. We took advantage of stem cells representing these lineages *in vitro* – ES (EPI) and XEN (PrE) cells – to have a scalable and tractable system for analysis. By establishing a bidirectional conversion system between XEN and ES cells by ectopic expression of either *Oct4*, *Klf4* and *Sox2* or of *Gata4*, we demonstrate for the first time a marked difference in the relative plasticity of the two lineages. While ES cells can be efficiently converted to iXEN cells, over ∼4 days and >95% efficiency, XEN cells reprogram to iPS cells with an extremely low efficiency, requiring ∼3 weeks and ∼0.2% efficiency. These differential kinetics and efficiencies are in line with the rare lineage switching events observed in early embryos (Chan et al., 2019; Nowotschin et al., 2019b; Xenopoulos et al., 2015).

The slow and inefficient reprogramming of XEN-to-iPS cells was notable for several reasons. First, OKS(M) reprogramming of numerous cell types has revealed an inverse correlation between the differentiation status and reprogramming amenability of a cell (Eminli et al., 2009; Tan et al., 2011). Similarly, transdifferentiation experiments suggest that cells with closer developmental relationships more readily interconvert (Graf and Enver, 2009; Vierbuchen and Wernig, 2011; Zhou and Melton, 2008). Therefore, XEN cells which represent an early embryonic sister lineage of EPI might be expected to have a relatively high potential for acquisition of an EPI/iPS state. In line with this reasoning, recent chemical reprogramming of somatic cells to iPS cells (or ‘ciPSCs’), suggested that a XEN-like state likely represents a plastic cell state which permits successful reprogramming to pluripotency (Guan et al., 2022; Li et al., 2017; Zhao et al., 2018, 2015). Moreover, XEN cells divide rapidly in culture, a feature that has been also associated with increased reprogramming efficiencies (Guo et al., 2014; Hanna et al., 2009). Despite these favorable properties, XEN cells showed a surprisingly low reprogramming potential, indicating an unusually stable state, which strongly resists acquisition of a pluripotent state. In support of this notion, scRNA-seq analysis revealed at least two major bottlenecks that XEN cells need to overcome to successfully reach the iPS state. The first bottleneck involves activation of key pluripotency genes, such as *Oct4* and *Rex1*, which is a common roadblock for most somatic cell types. The second bottleneck, requires downregulation of key XEN-associated genes, and appears unique to this cell type, since silencing of the somatic program is usually the first and most efficient milestone during reprogramming of most somatic cell types (Chronis et al., 2017; Polo et al., 2012; Sridharan et al., 2009; Stadtfeld et al., 2008). These observations highlight the unusual stability of the XEN/PrE program which cannot be easily overwritten by the potent Yamanaka factors (OCT4, SOX2, KLF4). This is partly due to a tight transcriptional network governed by the GATA factors, as perturbing GATA4 (but not GATA6) expression significantly increased XEN-to-iPS reprogramming efficiency. On the other hand, the “refractory” nature of XEN/PrE state could be due to a more rigid epigenetic/chromatin state that requires extensive remodeling to allow establishment of a new program. The latter is supported by our bulk and single-cell ATAC-seq data which detected a drastic and slow chromatin opening during XEN-to-iPS conversion, which could contribute to its low efficiency. The less favorable chromatin state of XEN compared to ES cells, especially around critical EPI and PrE gene loci, has been also reported before both at the level of DNA methylation and histone modifications (Bernstein et al., 2006; Rugg-Gunn et al., 2010; Senner et al., 2012). These findings suggest that the EPI and PrE fate bifurcation is accompanied by drastic epigenetic changes that support their differential developmental plasticity, thus posing a challenge for lineage conversion going from XEN to ES, but not vice versa.

In line with the contrasting dynamics of conversion and the epigenetic landscape of the two lineages, we noted that cells undergo somewhat divergent transcriptional changes during reprogramming in either direction, characterized by differential expression of chromatin modifiers and genes regulating cellular metabolism in the transition stages of lineage conversion. This finding partly reflects the epigenetic differences between the two lineages as noted previously, as well as metabolic differences that have been previously observed (Gatie and Kelly, 2018; Gatie et al., 2022; Mulvey et al., 2015). We also compared the *in vitro* trajectories to the EPI and PrE lineages *in vivo* from specification to differentiation and maturation. We find that the *in vitro* trajectories approximated sequential cell states in the embryo, mapping to the EPI and PrE lineages, but bypassing the uncommitted ICM progenitor state, indicating that this system is likely bistable, with the ICM perhaps representing an unstable *in vivo* state. This argues against models of tristability, and suggests that ICM lineage specification is a dynamic process with cells not occupying an uncommitted state in the absence of extracellular signals (Bessonnard et al., 2014; De Mot et al., 2016; Tosenberger et al., 2017). Notably, when comparing the *in vitro* with *in vivo* trajectories (**Figure 4F**), reprogramming XEN cells tracked along the PrE-like trajectory from the point of PrE lineage commitment throughout to upregulation of later PrE genes, such as *Gata4* and *Sox17*, in the blastocyst (Artus et al., 2011; Nowotschin et al., 2019b).

In conclusion, our studies clearly demonstrate the differential plasticity between ES and XEN cells and offer insights into the molecular basis of such differences. Further studies will likely determine the functional and developmental relevance of epigenetically restricting the PrE lineage soon after specification. A possible explanation is to restrict mixing of the embryonic and extraembryonic compartments to prevent “unfit” cells from disrupting development of the embryo-proper and establishment of the germ line. Extraembryonic cells tolerate polyploidy to a greater degree than cells of the embryo-proper, and also display unique genomic imprinting patterns (Eakin et al., 2005; Hudson et al., 2011; Ilgren, 1980; Takagi and Sasaki, 1975; Tarkowski et al., 1977). Therefore, ensuring that strict segregation of these two sister lineages may have evolved to ensure the survival, patterning and differentiation of tissues that constitute the embryo and the developments of the germline.

## ACKNOWLEDGEMENTS

We thank members of the Apostolou and Hadjantonakis labs for critical discussions and comments on the manuscript. We thank Christian Schroter MPI Dortmund, Germany for the Gata4 inducible mES cell line. Memorial Sloan Kettering Cancer Center’s (MSKCC) Mouse Genetics Core Facility (MGCF), Flow Cytometry Core Facility (FCCF), Integrated Genomics Operation (IGO) and the Single-Cell Analytics Innovation Lab (SAIL)) supported this work. MSKCC’s core facilities are funded by the NCI Cancer Center Support Grant (P30CA08748), with additionally funding for SAIL from the Alan and Sandra Gerry Center for Metastasis and Tumor Ecosystems, and the IGO from Cycle for Survival and the Marie-Josée and Henry R. Kravis Center for Molecular Oncology. This work was supported by an award from the STARR Tri-Institutional Stem Cell Initiative to AKH, EA and DP, with additional support from the NIH (R01DK127821, R01HD094868 and R01HD035455) to AKH, NIGMS (1R01GM138635) and the Tri-Institutional Stem cell Initiative (EA), and Howard Hughes Medical Institute to DP.

## AUTHOR CONTRIBUTIONS

V.G., E.A. and A.-K.H. conceptualized the study. V.G. generated embryo-derived stem cell lines, performed reprogramming, gene perturbations, flow cytometry, immunofluorescence and chimera experiments. V.G., S.N. and Y.-Y.K. generated single-cell genomics data. D.M and A.E.S. assisted with reprogramming experiments. A.P. performed analyses of bulk sequence data. Y.Y., M.S., R.S., and D.P. performed analyses of single-cell genomics data. V.G., E.A. and A.-K.H, drafted the manuscript, with input from all authors. A.-K.H. E.A. and D.P. acquired funding and supervised the work.

## MATERIALS AND METHODS

### Mouse strains and husbandry

All animal work was approved by Memorial Sloan Kettering Cancer Center’s Institutional Animal Care and Use Committee (protocol 03-12-017, Hadjantonakis PI). Animals were housed in a pathogen-free facility under a 12-hr light cycle. All embryos used for this study were obtained from natural matings of virgin females of 5-10 weeks of age. Mouse strains used in this study were: *Col1A1^TetO-OKSmCh/TetO-OKSmCh^*; *R26^M2rtTA/M2rtTA^* (Jackson Labs, Bar Harbor, ME, USA/stock ID: 034917) (Bar-Nur et al., 2014), *Oct4-GFP* (Jackson Labs, Bar Harbor, ME, USA/stock ID: 008214) (Lengner et al., 2007), and wild-type CD1 (Charles River). Genotyping PCR bands are as follows: *Col1A1^TetO-OKSmCh^*: wild-type – 331bp, knock-in – 551bp; *R26^M2rtTA^*: wild-type – 500bp, knock-in – 250bp; *Oct4-GFP*: wild-type – 434bp, knock-in – 234bp. Primers for genotyping are as follows:

### Cell lines, derivation and culture

XEN cell lines used in this study were: IM8A-1 (Kunath et al., 2005). Reprogrammable ‘3F’ XEN lines were derived from mice harboring *Col1a1^TetO-OKSmCh^*, *R26^M2rtTA^* and *Oct4-GFP* alleles using either TS cell conditions (3F4 line) or ES cell conditions (3F2, 3F6 and 3F9 lines) as detailed elsewhere (Niakan et al., 2013). Established XEN and iXEN cells were cultured in standard XEN cell culture conditions (Kunath et al., 2005; Niakan et al., 2013). Cells were seeded onto tissue culture grade plates coated with 0.1% gelatin (Millipore Sigma) for 5 mins at room temperature. Roswell Park Memorial Institute (RPMI) 1640 (Gibco) or Dulbecco’s modified Eagle’s medium (DMEM; Gibco) was supplemented with 15% FBS, 2 mM L-glutamine (Gibco), 1 mM sodium pyruvate (Gibco), 100 U/ml Penicillin, 100 µg/ml Streptomycin (Penicillin-Streptomycin; Gibco) and 0.1 mM 2-mercaptoethanol (Gibco).

ES cell lines used in this study were: R1 ES cells (Nagy et al., 1993) and *Col1a1^TetO-Gata4-mCherry/+^*;*R26^M2rtTA/+^*;*Gata6^H2B-Venus/+^*ES cells (Freyer et al., 2015; Schröter et al., 2015). ES cells and XEN-iPS cells were cultured in standard serum/LIF conditions as described previously (Czechanski et al., 2014). Cells were plated onto tissue culture grade plates coated with 0.1% gelatin for 5 mins at room temperature. DMEM was supplemented with 15% FBS, 0.1 mM non-essential amino acids (NEAA; Gibco), 2 mM L-glutamine, 1 mM sodium pyruvate, 100 U/ml Penicillin, 100 µg/ml Streptomycin, 0.1 mM 2-mercaptoethanol and 1000 U/ml leukemia inhibitory factor (LIF; prepared in house).

All cells were passaged every 2 days (∼80% confluence) by washing with phosphate buffered saline (PBS) followed by brief incubation in 0.05% Trypsin-EDTA (Gibco) at 37°C ∼2-3 mins). Trypsin activity was neutralized with serum-containing media (3x volume of Trypsin used) and dissociated cells were centrifuged at 400*g* for 3 mins before resuspending in culture media. Cells were replated at 1:8-1:10 dilution.

### XEN-to-iPS and ES-to-iXEN conversions

For XEN-to-iPS conversion, reprogrammable XEN cells were first cultured in standard XEN cell media with 2 µg/ml doxycycline (MP Biomedicals) for 2 days. mCherry-positive XEN cells were sorted using flow cytometry (see below) and plated onto mitomycin C-treated mitotically inactivated MEF feeders in 6-well or 10 cm plates for reprogramming. All plates were pre-gelatinized and layered with approximately 1×10^6^ MEF feeders per 50 cm^2^ culture surface area. mCherry-sorted XEN cells were reprogrammed in standard serum/LIF media supplemented with 2 µg/ml dox, 50 µg/ml ascorbic acid (Millipore Sigma) and 3 µM GSK3βi/CHIR99021 (Reprocell) (‘AGi’ media) (Bar-Nur et al., 2014). Media was replaced every two days. Doxycycline and ascorbic acid were prepared fresh every 5-7 days in dH_2_O and stored at 4°C protected from light. Doxycycline was filter sterilized using a 0.2 µm SFCA filter (Thermo Scientific). Reprogramming cells were dissociated with 0.25% Trypsin-EDTA prior to downstream analysis (Gibco).

ES-to-iXEN conversion was carried out as described previously (Wamaitha et al., 2015). ES cells were plated onto pre-gelatinized plates in standard serum/LIF media supplemented with 2 µg/ml dox. Media was replaced every two days. Cells were dissociated with 0.05% Trypsin-EDTA prior to downstream analysis.

XEN and ES cells were plated at the following densities for reprogramming:

| ***Format*** | ***Culture days*** | ***XEN Number*** | ***ES Number*** |
| --- | --- | --- | --- |
| 6-well | 1 | --- | 200-250K/well |
| 6-well | 2 | 150-200K/well | 150-200K/well |
| 6-well | 3 | --- | 100-150K/well |
| 6-well | 4 | 100-120K/well | 40-50K/well |
| 6-well | 5 | --- | 8-10K/well |
| 6-well | 6 | 20-30K/well | --- |
| 6-well | 8 | 5-8K/well | --- |
| 6-well | 10 | 3-4K/well | --- |
| 6-well | 12 | 1-1.5K/well | --- |
| 6-well | 14 | 500-1K/well | --- |
| 6-well | 16 | 500-1K/well | --- |
| 10 cm | 7 | 50-75K | --- |
| 10 cm | 14 | 5-10K | --- |

### iPS-to-iXEN conversion

a *pCX-Gata4-E2A-E2-Crimson* expression vector was first generated using a *pCAGGS* vector (Hitoshi et al., 1991) as a backbone digested with EcoRI restriction enzyme. Coding sequences for mouse *Gata4* and *E2-Crimson* preceded by an E2A self-cleaving peptide (*E2A-E2-Crimson*) were assembled using NEBuilder HiFi DNA assembly (NEB) to generate *pCX-Gata4-E2A-E2-Crimson*. iPS cells derived from ProB cells from mice harboring *Col1a1^TetO-OKSmCh^*, *R26^M2rtTA^* and *Oct4-GFP* alleles were transfected with *pCX-Gata4-E2A-E2-Crimson* plasmid using Lipofectamine 3000 (Invitrogen). *Oct4*-GFP+/PDGFRα+ and Oct4-GFP-/ PDGFRα+ cells were flow sorted on Day 5 or Day 7 of conversion, then plated onto XEN-to-iPS reprogramming conditions and analyzed for marker expression 14 days following sorting and plating. In parallel, freshly converted iXEN cells (passage 2 following iXEN formation) from ProB iPS cells were also plated and analyzed for marker expression by flow cytometry.

### Immunophenotyping by flow cytometry and fluorescence activated cell sorting

Dissociated cells were filtered through a 0.35 µm nylon mesh strainer (Falcon) to achieve a single cell suspension. Cells were centrifuged at 400*g* for 5 mins at room temperature and washed twice with wash buffer (5% FBS in PBS without Ca^2++^ and Mg^2++^). Cell pellets were then resuspended in 100 µl staining buffer (wash buffer with diluted antibodies) and incubated at room temperature for 30 mins protected from light. Following incubation, cells were washed twice with wash buffer. Cell pellets were finally resuspended in wash buffer containing 1 µg/ml 4′,6-diamidino-2-phenylindole (DAPI; Invitrogen) or 7-AAD viability dye (BioLegend; 1 µl/sample). The following antibodies were used per 100 µl of staining volume for 1×10^6^ cells: PE-Cy7::CD140a (PDGFRα; eBioscience) at 0.0625 µg/µl, AlexaFluor647::SSEA-1 (BioLegend) at 2.5 µl/sample or BV421::SSEA-1 (BioLegend) at 2.5 µl/sample. Stained cell samples were then analyzed for marker expression using LSRFortessa (BD Biosciences). DAPI and BV421 were excited at 405 nm and detected using 450/50 nm band-pass filters. GFP and Venus were excited at 488 nm and detected using 525/50 nm band-pass filter. mCherry was excited at 561 nm and detected using 610/20 nm band-pass filter. 7-AAD was excited at 561 nm and detected using 670/30 nm band-pass filter. PE-Cy7 was excited at 561 nm and detected using 780/60 nm band-pass filter. AlexaFluor647 was excited at 633 nm and detected using 670/30 nm band-pass filter. For cell sorting, samples were prepared as described and resuspended at a final concentration of 5-10×10^6^/ml in wash buffer. Cells were sorted using SORP FACSAria IIu (BD Biosciences) with a 100 µm nozzle at 20 psi. Gating strategies are provided in **Figure S2A**. Data were analyzed using FlowJo and R (http://www.r-project.org/).

### Real-time quantitative PCR

Cells were harvested following trysinization and washed once with PBS. Total RNA was extracted from cells pellets using TRIzol reagent (Invitrogen) according to manufacturer’s directions. cDNA synthesis was carried out with 1 µg of RNA using the QuantiTect Reverse Transcription Kit (Qiagen) and subsequently diluted 1:25 in dH_2_O. 5 µl of resulting cDNA was combined with 1 µM each of forward and reverse primers and 10 µl of PowerUp SYBR Green Master Mix (Applied Biosystems) for RT-qPCR in 20 µl of total volume. RT-qPCR reactions were carried out in a CFX96 Real-Time PCR detection system (BioRad). All analyses were carried out in R. Raw Ct values of three technical replicates per reaction were averaged and normalized to the mean of two reference genes: *Actb* and *Gapdh*. Normalized Ct values were then plotted as “expression” values using the following equation: *y = 2^-Ct^*. The following primers were used:

### Immunofluorescence

#### Cultured cells

Cells were washed twice for 5 mins. each with PBS before fixation at room temperature with 4% PFA (Electron Microscopy Sciences) for 15 mins. Fixed cells were washed again with PBS twice for 5 mins. each before permeabilization with PBST [PBS + 0.1% Triton-X 100 (Millipore Sigma)] at room temperature for 10 mins. Cells were subsequently blocked with blocking buffer [PBST + 1% bovine serum albumin (BSA; Millipore Sigma) + 3% donkey serum (Jackson ImmunoResearch)] at room temperature for 15 mins. Cells were then incubated with primary antibodies diluted in blocking buffer overnight at 4°C. Following incubation, cells were washed three times for 10 mins. each with PBST. Cells were then incubated for 2 hrs. with secondary antibodies diluted in blocking buffer at room temperature. Next, cells were washed twice for 10 mins. each with PBST before incubation with DAPI diluted 1:10,000 in PBST for 10-15 mins. DAPI was washed off with PBS before imaging.

#### Post-implantation stage embryos

Freshly dissected embryos were fixed in 4% PFA at room temperature for 15 mins. Following fixation, embryos were washed once with PBST before permeabilization in 0.5% Triton-X (in PBS) at room temperature for 30 mins. Fixed and permeabilized embryos were then blocked with blocking buffer (PBST + 1% BSA + 5% donkey serum) overnight at 4°C. Next, embryos were incubated with primary antibodies diluted in PBST + 1% BSA overnight at 4°C. The following day, embryos were washed 3 times for 10 mins. each in PBST. Embryos were then blocked for 5 hrs. at room temperature prior to incubation with secondary antibodies diluted in blocking buffer overnight at 4°C. Stained embryos were then washed twice with PBST for 10 mins. each before incubation with DAPI diluted 1:1000 in PBST. Embryos were finally washed in PBS before imaging.

Secondary AlexaFluor antibodies (donkey/IgG; Invitrogen) were used at 1:500 dilution. See Key Resources Table for list of primary antibodies used in this study.

### XEN-iPS cell-embryo chimeras

Prior to generating chimeric embryos, XEN and XEN-iPS cells were labeled with an *mCherry* expression vector. For this, a *pCX-mCherry* expression vector (unpublished) was used in which *mCherry* is inserted into a *pCAGGS* vector (Hitoshi et al., 1991) as previously described (Nowotschin et al., 2009; Okabe et al., 1997). The plasmid was linearized by *ScaI* restriction digest and purified using the QIAquick PCR purification kit (Qiagen). Purified linear plasmid was transfected using Lipofectamine 3000 (Invitrogen). Stably expressing mCherry-positive XEN cells were flow sorted, and mCherry-positive XEN-iPS clones were picked 10 days following transfection and expanded prior to 8-cell morula injection.

XEN-iPS chimera embryos were generated by the Mouse Genetics Core at MSKCC. XEN or XEN-iPS cells were injected into C57Bl/6 host 8-cell morula, and transferred to pseudo-pregnant females. Chimeric embryos were recovered at E5.5-E6.0.

### Genetic editing of XEN cells

To knockout *Gata4* or *Gata6* gene expression in XEN cells, the PX458 vector (Addgene #48138) was first modified to express E2-Crimson instead of EGFP. The resulting PX458-E2-Crimson vector was digested using BbsI-Hf (NEB) and single guide RNA (sgRNA) targeting *Gata4* or *Gata6* was annealed as previously described (Ran et al., 2013). Top 3 ranked sgRNAs were tested from the CHOPCHOP tool (Labun et al., 2019). Assembled Cas9/sgRNA plasmids were transfected into XEN cells using Lipofectamine 3000 (Invitrogen). E2-Crimson-positive cells were assessed for successful knockout of gene expression and reprogramming potential (see **Figure 5B-C** and **S6C**). pSpCas9(BB)-2A-GFP (PX458) was a gift from Feng Zhang (Addgene plasmid # 48138; http://n2t.net/addgene:48138; RRID:Addgene_48138).

### Image acquisition and processing

Brightfield and epifluorescence images shown in **Figure 1A-B** were acquired on a Zeiss Axio Vert.A1 inverted microscope with a black and white camera (Axiocam MRm). Brightfield and epifluorescence images shown in **Figure S1E** were acquired on a Zeiss AxioZoom stereomicroscope with a Zeiss Axiocam MRc CCD camera and ZEN 2.3 software, using the manual extended depth of focus application. Immunofluorescence images shown in **Figure 1E**, **S1B** and **S6C** were acquired on a Zeiss LSM 880 laser-scanning confocal microscope. Post-implantation stage embryos were imaged in a microdrop of PBS on a 35 mm glass bottom dish (MatTek) using a Plan-Apo 20×/NA0.8 M27 objective. Z-stacks were taken at 0.88-μm intervals. Images in **Figure S1B** and **S6C** were acquired using an EC Plan-Neofluar 40×/NA1.30 oil immersion objective at 1-μm z-intervals. Fluorescence was excited using a 405-nm diode (Hoechst 3342), 488-nm argon, 561-nm DPSS-561-10 and HeNe 633-nm lasers. Raw image data were processed using ImageJ (Rasband, W.S., ImageJ, U.S. National Institutes of Health, Bethesda, Maryland, USA, https://imagej.nih.gov/ij/, 1997–2018). Raw images for **Figure S6C** were processed using Imaris (Bitplane), where nuclei were identified using the Spot model, and corresponding relative fluorescent intensities were measured for each channel. Data were analyzed using R (http://www.r-project.org/).

### Bulk RNA-seq analysis

Paired-end sequenced reads from ES, XEN and XEN-iPS cell lines were aligned to mouse genome (mm10) with Tophat2 (version 2.1.1) with default setting and “-r 200 –mate-std-dev 100” option. Sorting of aligned reads was performed with samtools (Li et al., 2009) and reads were assigned to protein coding and long-non coding genes (Mus_musculus.GRCm38.95.gtf) with the use of htseq-count (Anders et al., 2015) and ‘-m intersection-nonempty’ option. DESeq R package (Anders and Huber, 2010) was used to call differentially expressed genes between XEN and ES cell lines. Only genes with p-adj (<0.01) and fold change cut off (2) were considered as differentially expressed between ES and XEN.

### Sample preparation for single-cell RNA-seq and single-cell ATAC-seq

XEN-to-ES and ES-to-XEN conversions were carried out as described for the durations outlined in **Figure 3A** prior to collection. For ‘starting’ populations, XEN cells were cultured on MEF feeders for 7 days in reprogramming media without dox induction (i.e., AGi media minus dox), and ES cells were cultured in standard serum/LIF conditions prior to collection. To collect nascent XEN-iPS cells, XEN cells were reprogrammed for 14 days, sorted for SSEA-1-GFP+ PDGFRα-cells using flow cytometry (see above) and replated for an additional 14 days (day 28 of reprogramming). At this point SSEA-1+ GFP+ PDGFRα-cells were sorted and replated onto MEF feeders in standard serum/LIF media for 7 days prior to dissociation. A similar strategy was applied for nascent iXEN cells – ES cells were treated with dox for a period of 4 days, by when >95% of cells have successfully converted to iXEN (see **Figure 2**). Following this period, dox was withdrawn from the culture media for 7 days prior to collection. As described above and schematized in **Figure S3A**, a cell sorting strategy was used for XEN-to-ES converted day 7, day 14 and day 28 cells, and ES-to-XEN converted 9h cells. ES-to-XEN 24h and 48h samples were not sorted.

All cells were dissociated using either 0.05% or 0.25% Trypsin-EDTA and dissociated into single cells by repeated pipetting and passing through a 0.35 µm nylon mesh strainer prior to downstream processing. For scRNA-seq samples, dissociated or sorted cells were counted and diluted in DMEM + 10% FBS. Final cell suspensions were loaded on a Chromium Controller targeting a 5,000 – 8,000 cell range, depending on the sample, to generate single-cell 3’ RNA-seq libraries in duplicate (Zheng et al., 2017). For scATAC-seq samples, cells were washed in PBS + 0.04% BSA and processed for nuclei isolation following manufacturer’s instructions. Final nuclei preparations were loaded on a Chromium Controller targeting 10,000 nuclei, and processed in singlicate. Libraries were generated following the manufacturer’s instructions (10x Genomics Chromium Single Cell 3′ Reagent Kit User Guide v2 Chemistry and 10x Genomics Chromium Next GEN Single Cell ATAC Reagent Kit Guide v1.1).

To obtain single parietal endoderm cells from E7.5 and E8.5 wild-type mouse embryos, we first dissected the embryo out of its decidua, and then teased the parietal yolk sac out the decidua in DMEM/F12, 5% Newborn Calf Serum (Gibco). To remove debris, the tissue was washed five times in 200µl drops of DMEM/F12, 5% Newborn Calf Serum shaking, followed by five washes in DMEM/F12. The parietal yolk sac was then incubated in 100 µl TrypLE (Invitrogen) for 15 min at room temperature for dissociation into single cells. For the subsequent mechanical dissociation 100 µl DMEM/F12, 20% Newborn Calf Serum, 4 mM EDTA was added. The parietal cell clumps were dissociated into single cells first by using a P200 pipette to shake off Reichert’s membrane followed by mouth pipetting with pulled glass capillaries. The resulting single cell suspension was filtered through a FlowMi cell strainers (4 µm, Millipore Sigma) to remove cell clumps and debris and centrifuged, and pellet resuspended into DMEM/F12, 10% Newborn Calf Serum. Cells were stored on ice until loaded onto a Chromium Controller (10x Genomics) targeting 3,000 - 4,000 cells (E7.5), and 9,000 cells (E8.5) to generate single-cell 3’ RNA-seq libraries in duplicate. Of note, our efforts to isolate single parietal endoderm cells from Reichert’s membrane of earlier staged (E5.5 and E6.5) embryos yielded too few cells to load onto a 10x Genomics Chromium chip.

### Next-generation sequencing of single-cell libraries

Single-cell 3′ RNA-seq and single-cell ATAC-seq libraries were quantified on an Agilent Bioanalyzer with a high-sensitivity chip (Agilent), and Kapa DNA quantification kit for Illumina platforms (Roche). Libraries were pooled according to target cell number loaded for a sequencing depth of 20K-25K (for scRNA-seq) or 35K (for scATAC-seq) reads per cell and accounting for the capacity of an Illumina NovaSeq flow cell. Library pools were loaded on an Illumina NovaSeq 6000 using 2× NovaSeq 6000 S2 reagent kits (200 cycles) and 1× NovaSeq 6000 S4 reagent kits (300 cycles) using the following read length: 26-bp read 1, 8-bp I7 index and 98-bp read 2 (for scRNA-seq), or 50-bp read 1, 8-bp I7 index, 16-bp I5 index and 50-bp read 2 (for scATAC-seq). Libraries of ParE cells isolated from E7.5 and E8.5 embryos were sequenced as previously published (Nowotschin et al., 2019b).

### Single-cell RNA-seq data processing

#### Data preprocessing

scRNA-seq data from each sample was preprocessed using the SEQC pipeline (Azizi et al., 2018) using GRCm38/mm10 mouse genome and default SEQC parameters to obtain molecule count matrices. The SEQC pipeline aligns the reads to the genome, corrects barcode and unique molecular identifier (UMI) errors, resolves multi-mapping reads, and generates a molecule count matrix. SEQC also performs a number of filtering steps: (1) Identification of true cells from cumulative distribution of molecule counts per barcode, (2) removal of apoptotic cells identified at cells with >20% of molecules derived from the mitochondria, and (3) removal of low-complexity cells identified as cells where the detected molecules are aligned to a small subset of genes. The filtered count matrix was normalized by dividing the counts of each cell by the total molecule counts detected in that particular cell. The normalized matrix was multiplied by the median of total molecules across cells to avoid numerical issues. Normalized data were log transformed with a pseudo-count of 0.1.

#### Data clean-up, dimensionality reduction and visualization

Each reprogramming trajectory was analyzed separately at first by pooling the two replicates. For each trajectory, 1500 highly variable genes were selected using scanpy (Wolf et al., 2018). Data was projected onto principal components following gene selection to overcome the noise in scRNA-seq data due to high degree of dropouts. The number of components that explain 85% of the variance were retained for each of the downstream analyses. Force directed layouts were computed for each trajectory by first computing an adaptive kernel (van Dijk et al., 2018) to account for large density differences in the data, computed using the Palantir package (Setty et al., 2019) with default parameters. For the XEN-to-iPS dataset, a cluster of cells with fibroblast signature and another with 2-cell stage signature were removed. For the ES-to-iXEN dataset, a cluster of spontaneously differentiating cells were removed from the starting ES cell samples.

#### Batch correction

We did not observe any batch effects amongst the replicates of the XEN-to-iPS dataset and all conditions and replicates from XEN-to-iPS dataset were pooled for downstream analysis. On the other hand, we did observe batch effects in ES-to-iXEN timepoints. Batch effect correction was performed using mnnCorrect (Haghverdi et al., 2018) using replicate 1 as the reference. The pooled datasets were analyzed using the same procedure described above with one difference – 2500 highly variable genes were used to account for greater heterogeneity of the data across timepoints. The pooled ES-to-iXEN results demonstrate that batch effects were corrected effectively.

### Trajectory analysis

Palantir was used for analysis of the XEN-to-iPS and ES-to-iXEN conversion trajectories using default parameters (Setty et al., 2019). Palantir models differentiation as a Markov chain and computes for each cell the probability of differentiating to each of the terminal states of the system. The terminal states are also determined automatically by Palantir. Terminal states were specified for the *in vivo* trajectories since end points were known. Palantir first computes diffusion maps, a low dimensional representation of the phenotypic space occupied by the cells. A nearest neighbor graph is then constructed in the diffusion map space. Shortest path distances through this graph from a pre-designated start cell is used to determine a pseudotime ordering of cells. Pseudotime order is then used to transform the nearest neighbor graph into a Markov chain based on which the branch probabilities are computed. The branch probabilities for each cell are summarized using entropy to compute the differentiation potential, a predicted measure of plasticity of the cells.

#### Diffusion maps and MAGIC imputation

To compute diffusion maps (Coifman and Lafon, 2006; Haghverdi et al., 2015; Setty et al., 2016) A k-nearest neighbor graph (*k* = 50) was constructed using Euclidean distance with principal components as inputs. The distance matrix representing this graph was converted to an affinity matrix using the adaptive anisotropic kernel i.e., for each cell, distance to *l*^th^ neighbor (*l* (17) < *k* (50)) was used as the scaling factor to account for differences in densities in data. The affinity matrix was normalized to generate the diffusion operator. The top Eigenvectors from the Eigenvalue decomposition of this operator, termed diffusion components, represent the low-dimensional embedding of the data. The number of components was chosen by the Eigen gap among the top Eigen vectors. The same diffusion operator was used for MAGIC (van Dijk et al., 2018) imputation of gene expression data (for t = 3 steps). Single-cell gene expression plots throughout the manuscript use MAGIC imputed expression. Diffusion component computation and MAGIC imputation were performed using the Palantir package (https://github.com/dpeerlab/Palantir).

#### Application of Palantir to characterize transdifferentiation trajectories

XEN-to-iPS and ES-to-iXEN scRNA-seq data were analyzed separately. A random cell from timepoint 0 was used as the input start cell, which adjusts the start to the nearest extreme of the diffusion components. Palantir automatically determined the terminal states in each trajectory, including a single terminal state in the ES-to-iXEN trajectory, representing the final iXEN state. By contrast, three states – T1, T2 and T3 (final iPS state) – were identified in the XEN-to-iPS trajectory.

#### Comparing XEN-to-iPS reprograming terminal states by differential gene expression analysis

We sought to identify the genes that are enriched in each of the terminal (and start) states of the XEN-to-iPS trajectory (i.e., Start, T1, T2, T3). For this, we identified the XEN clusters that contained these terminal points (Cluster 2 – XEN/Start, Cluster 4 – T3/iPS, Cluster 7 – T1 and Cluster 11 – T2) and computed the genes that are differentially expressed in each of these clusters compared to rest of the cells in the trajectory using MAST (Finak et al., 2015). To summarize the results, we collected the top 50 genes that are significantly differentially expressed (FDR adjusted p-value < 0.01 and logFoldChange > 2) in each of these clusters. We then took the union of all the obtained list of genes and displayed the average z-scored expression of these genes in each of the terminal points as a heatmap using clustermap function in the Seaborn package (**Figure S3B**).

### Comparison of the two reprogramming trajectories

To enable a direct comparison between the XEN-to-iPS and ES-to-iXEN transitions, we co-embedded the two trajectories using Harmony (Nowotschin et al., 2019b). Harmony augments the nearest neighbor graph within each trajectory (*k* = 50) with mutual nearest neighbors (*k* = 50) between XEN-to-iPS and ES-to-iXEN conversions, and uses them to estimate a joint affinity matrix. The nearest neighbor distance matrix is converted into an affinity matrix using an adaptive Gaussian kernel, where for each cell the affinity is defined as the negative exponential of the distance to a neighbor, scaled by the distance to it’s *l*^th^ neighbor (*l* = *k*/3 = 17). Once this joint affinity matrix is constructed, it is normalized to obtain a Markov matrix, which is then used as an input to the Force Directed Layout embedding as implemented in the Harmony package.

#### Comparison of in vitro trajectories during intermediate transition phase

Based on the embedding on the FDL (**Figure 4A**), we reasoned that while the terminal points of the XEN-to-iPS and ES-to-iXEN trajectories are phenotypically similar, the cells at the transition stages appear distinct. We therefore sought to investigate if XEN-to-iPS and ES-to-iXEN reprogramming follow the same phenotype trajectories. For this, we assumed a batch effect as a source of this discrepancy between the two trajectories and began by aligning them using Harmony (default parameters as implemented in Scanpy.external package in Python) (Korsunsky et al., 2019). We then performed PhenoGraph clustering as implemented in Scanpy.external package in Python (clustering_algo = leiden, *k* = 30, resolution_parameter = 3) to obtain 29 clusters. From the obtained clusters, we identified those consisting of cells in the transition state (**Figure S5B**). Among these selected clusters, we computed significantly differential genes between cells transitioning from XEN-to-iPS and ES-to-iXEN using MAST with default parameters (Finak et al., 2015), followed by GSEA analysis using GO annotations (http://www.gsea-msigdb.org/gsea/msigdb/human/genesets.jsp?collection=C5).

To ensure our inference is not impacted by the choice of batch correction method, we used an alternative clustering strategy. In particular, we used Harmony (Nowotschin et al., 2019b) and grouped cells using Spectral Clustering (K-means with n_clusters = 30 as implemented in sklearn package in Python) on the computed diffusion components (top 14 eigenvectors of the joint Markov matrix identified based on the eigengap). We repeated the computation of differentially expressed genes and enriched gene sets and obtained highly similar results.

#### Comparison of in vivo and in vitro trajectories

The goal of this analysis is to map the path taken by cells in the *in vitro* reprogramming trajectories onto the *in vivo* developmental trajectories to assess whether the *in vivo* developmental states are recapitulated in the *in vitro* trajectories.

#### Construction of the in vivo trajectory

scRNA-seq data from mouse embryos at the following stages were used for constructing the *in vivo* trajectory: E3.5, E4.5, E5.5, E7.5, E8.75 using ICM, epiblast, primitive & visceral endoderm and parietal endoderm cells (Nowotschin et al., 2019b). Since the data was generated at discrete timepoints, we used our Harmony algorithm (Nowotschin et al., 2019b) to connect successive timepoints. Harmony augments affinity matrix derived from the nearest neighbor graph with mutually nearest neighbors between successive time points. An affinity matrix is derived from this augmented nearest neighbor graph which serves as input for downstream trajectory analysis and imputation. In addition to using mutually nearest neighbors between successive time points, we also utilized mutually nearest neighbors between E4.5 and E7.5 parietal endoderm cells since parietal endoderm cells were not captured from E5.5 and E6.5 in the *in vivo* dataset, and between E7.5 and E8.75 parietal endoderm cells. The augmented affinity matrix was used as input to compute diffusion components and Palantir was used to compute a pseudotime ordering using ICM as the start and epiblast, parietal and visceral endoderm cell states as the terminal state inputs. More specifically, Harmony was applied on the joint PCA embedding (>85% of variance explained) and *n_neighbors = 30*, and Force Directed Layout using *sc.tl.draw_graph* function in Scanpy was run to visualize the augmented embedding.

#### Mapping between in vivo and in vitro trajectories

Comparison between *in vivo* and *in vitro* trajectories were undertaken separately for XEN-to-iPS and ES-to-iXEN datasets. Following the construction of the *in vitro* trajectory using Palantir and *in vivo* trajectory using Harmony and Palantir, we mapped the two to compare the path taken by *in vitro* cells. First a cell-by-cell affinity matrix was derived using the nearest neighbor graph constructed using all *in vivo* and *in vitro* cells. This affinity matrix was augmented with mutually nearest neighbors between *in vivo* timepoints to recapitulate the *in vivo* trajectory using Harmony (Nowotschin et al., 2019b). Harmony was again used to then add mutually neighboring edges between *in vivo* and *in vitro* cells. Thus, the final augmented affinity matrix comprised of the following set of edges: (i) nearest neighbor edges across all cells, (ii) edges between successive *in vivo* timepoints and (iii) edges between *in vivo* and *in vitro* cells. Harmony was applied using default parameters. Highly variable genes (1500) from the *in vivo* trajectory were used for this analysis.

The augmented affinity matrix served as input to compute diffusion maps using Palantir with default parameters. These diffusion components contain information about the *in vivo* and *in vitro* trajectories and the relationship between them, making the comparison feasible. Rather than compare individual cells, we binned the two trajectories into equal sized bins using their respective pseudotime order (**Figure S4**). For the *in vivo* trajectory, bins were separated based on lineage. Cells were binned using 20 intervals. Each *in vitro* bin was mapped to its closest *in vivo* bin using the mean multi-scale distance (Setty et al., 2019) between each pair of *in vivo* and *in vitro* cells in the augmented *in vivo*-*in vitro* diffusion space. The median position of the cells in the nearest *in vivo* bin was used for representing *in vitro* bins in **Figure 4F** and **S5E-F**.

### Clustering of gene expression trends

XEN-to-iPS cells were first clustered using PhenoGraph (Levine et al., 2015) (*k* = 30) using the principal components as inputs. Differentially expressed genes were identified in each cluster using MAST (Finak et al., 2015) with p-value < 1e-5 and log fold change > 1.5. For each cluster, cells from all other clusters were used as a baseline for comparison. Gene expression trends along XEN-to-iPS pseudotime order for each differentially expressed gene were computed using the gene trend analysis as implemented within the Palantir package, which in turn utilizes the Generalized Additive Models (GAMs; gam package in R) (Hastie and Tibshirani, 1990). These gene expression trends were then z-scored and used to cluster the genes. More specifically, we first computed a gene-gene nearest neighbor graph using NearestNeighbor function in sklearn package in Python using “radius = 0.025” and “metric = ‘correlation’” parameters. The distance matrix was then symmetrized and converted into an affinity matrix defined as 1-distance. We then ran Louvain clustering algorithm on the obtained affinity matrix. Finally, we excluded any clusters with only one gene (i.e., singleton clusters) to obtain a final set of clusters of genes with similar gene trends.

### Bulk ATAC-seq analysis

Paired-end sequenced reads from replicate ES and XEN cells were aligned to mouse genome (mm10) with Bowtie2 (version 2.3.4.1) (Langmead and Salzberg, 2012) and “--local –very-sensitive-local-I 10 × 2000” option active. Alignment was followed by filtering of low quality reds (MAPQ<20), duplicate reads, chrM reads and blacklisted regions with the use of Samtools (Li et al., 2009), “MarkDuplicates” from picard tools and bedtools (Quinlan and Hall, 2010). All filtered reads were corrected for Tn5 insertion at each read end by shifting +4/-5 bp from the positive and negative strand, respectively, and peak calling was performed with MACS2 (version 2.1.1) (Zhang et al., 2008) ‘—narrow’ option active and default settings. Non-overlapping peaks from replicates were filtered out and only common peaks were used. Peak center (summit file) generated with MACS2 with ‘—narrow’ option was extended to 100bp (+/-50bp) for motif search and all overlapping summits were merged to form an accessibility atlas which was used as background for motif and ChIP enrichment with LOLA R package (see below for LOLA enrichment analysis). For measuring relative accessibility of genes enriched at XEN-to-iPS terminal states, signal from all accessible regions around the promoter regions of selected genes (scRNA-seq gene lists/groups) were summed and compared with the use of Mann-Whitney U test.

### Single-cell ATAC-seq data processing

#### Data processing and metacell analysis

Cell Ranger ATAC (Satpathy et al., 2019) was used to preprocess the scATAC-seq data based on GRCm38/mm10 mouse genome, to obtain the sequence alignment files and fragment files, which are then provided as input to the ArchR software (Granja et al., 2021). Cell Ranger ATAC performs barcode location detection, sequencing error correction, read alignment, and duplicate read pair identification. The resulting fragment file contains the genomic position information of each sequenced scATAC-seq fragment and the identity of the corresponding cell. With the preprocessed scATAC-seq data in each of the two conversion trajectories (XEN-to-iPS and ES-to-iXEN), ArchR was used to identify chromatin accessibility peak loci from the data. Each chromatin accessibility peak locus corresponds to an accessible genomic region. Specifically, ArchR uses the input files to generate a sparse count matrix where each row corresponds to a single cell and each column corresponds to a genomic bin (500 base pair). The values in the matrix are the number of fragments in each genomic bin of each cell. ArchR employs the iterative Latent Semantic Indexing (LSI) approach (Granja et al., 2019; Satpathy et al., 2019) to perform normalization of the sparse count matrix using the frequency-inverse document frequency (TF-IDF) method (Cusanovich et al., 2015). We used 100K as the number of variable features for the LSI implementation with the other parameters as default. Next, singular value decomposition (SVD) is applied to the normalized count matrix for dimension reduction of the scATAC-seq data. We used 30 as the number of dimensions after reduction. ArchR then performs clustering of the cells using the graph clustering approach from Seurat (Hao et al., 2021) and generates pseudo-bulk replicates based on the cell groups identified from the clustering. More specifically, the data of a set of single cells sharing similarity are merged to create a pseudo-sample to address the sparsity problem of scATAC-seq data. Peak calling was performed using MACS2 (Zhang et al., 2008) on the pseudo-bulk replicates and the iterative overlap peak merging procedure (Corces et al., 2017) was applied in ArchR to generate a merged set of chromatin accessibility peak loci for the single cells. Each peak locus is 501bp in length.

The XEN-to-iPS trajectory consists of five time points: start, day 7, day 14, day 28, and end (sorted iPS cells), with one replicate for each time point. The ES-to-iXEN trajectory consists of five time points: start, 9h (h: hour), 24h, 48h, and end (iXEN cells), with two replicates for 24h and 48h and one replicate for each of the other time points. The replicates were merged for chromatin accessibility peak loci identification in each trajectory. There are 61040 and 95396 single cells in the scATAC-seq data of the XEN-to-iPS and ES-to-iXEN trajectories, respectively. 257618 and 250671 chromatin accessibility peak loci were identified in either trajectory using ArchR, respectively.

Next, the SEACells algorithm (Persad et al., 2022) was used to identify metacells, each of which are representative of a small assembly of single cells sharing the same or similar cell states based on the scATAC-seq data in each conversion trajectory. We identified 543 metacells in the XEN-to-iPS trajectory and 567 metacells in the ES-to-iXEN trajectory (default parameters, except *n_waypoint_eigs = 10*, *waypoint_proportion = 1*). Each metacell is associated with 112 single cells on average or 168 single cells on average in the XEN-to-iPS and ES-to-iXEN trajectories, respectively. The scRNA-seq read counts for each gene in each metacell is computed as the summed expression over all the single cells assigned to that metacell. The scATAC-seq read counts in each peak locus across the single cells represented by the same metacell were aggregated to approximate the accessibility of the locus in the corresponding metacell, to overcome the limitation of sparsity in scATAC-seq data. The chromatin accessibility count matrix of the scATAC-seq metacells were then normalized – the read count in each peak locus in each metacell was divided by the total read count in the metacell and multiplied by the median of the total counts per metacell across metacells. We performed log transformation of the normalized count matrix with a pseudo-count of 1. The subsequent scATAC-seq data analyses were conducted at the metacell level.

Next, we employed the highly variable gene identification function in the Scanpy package (Wolf et al., 2018) to detect the peak loci with highly variable accessibility (denoted as highly variable peak loci) from the scATAC-seq data of the metacells. Specifically, dispersion of the chromatin accessibility across the metacells was calculated for each peak locus and normalized within each group of peak loci sharing similar mean accessibility across the metacells. The peak loci with normalized accessibility dispersion and mean accessibility larger than the specified thresholds were selected as a set of highly variable peak loci and were used for representation of the chromatin accessibilities in the metacells in either trajectory, in order to capture more distinctive features across different cell states. We chose a threshold of 3.0 for the normalized accessibility dispersion and a threshold of 0.0125 for mean accessibility in both conversion trajectories. We identified 3584 and 1955 highly variable peak loci with the specified thresholds in the XEN-to-iPS and ES-to-iXEN trajectories, respectively.

For both conversion trajectories, we performed principal component analysis (PCA) (Jolliffe and Cadima, 2016) for dimension reduction of normalized and log-transformed chromatin accessibility matrix of highly variable peak loci for the metacells and selected the first 100 principal components (PCs) for feature representation. The force-directed layout (FDL) plots of metacells were generated using the Palantir package (Setty et al., 2019) with default parameters for visualization. For the XEN-to-iPS trajectory, PhenoGraph clustering (Levine et al., 2015) was applied to the metacells using default parameters (*k* = 30) and the feature representation from highly variable peak loci, identifying 9 clusters (**Figure 6B**). We further assigned the clusters of metacells to 7 major groups based on the time points from which they were collected, and their relative accessibility patterns along pseudotime from the XEN state to the iPS cell state. Specifically, group 1 mostly comprises metacells from the start time point. Group 2A and 2B correspond to the two clusters identified for metacells from day 7. Group 3 and 4 (4A and 4B) each contain a mixture of the metacells from day 14 and day 28. The members in group 4B are relatively closer to the iPS cell state while group 4A are more dispersed in the cell states. Group 5 predominantly contains metacells in the iPS cell state. In the ES-to-iXEN trajectory, the cell groups are distinguishable by the associated time points, (**Figure 6D**), and metacells were thus clustered by time point.

### TF binding activity estimation from scATAC-seq data

We used the chromVAR method (Schep et al., 2017) to estimate transcription factor (TF) binding activities in each metacell. For each metacell and each TF with binding motifs, chromVAR aggregates the accessibility of the peak loci where the binding motif of the TF is identified, and computes a bias-corrected z-score of the aggregated accessibility as the TF binding activity score in the metacell (noted as chromVAR score). Specifically, the original z-score measures the deviation of the aggregated accessibility for the TF in a metacell from the mean value of the aggregated accessibility across the metacells. For each peak locus containing the TF binding motif, peak loci with GC content and mean chromatin accessibility across the metacells both matching the corresponding locus were sampled genome-wide, forming sets of background peak loci to estimate the background distribution of the z-score for bias-correction. To identify TF binding motifs in the peak loci, we used the curated CIS-BP mouse TF binding motif collection retrieved from the chromVAR repository (Schep et al., 2017) and the matchMotifs function in the motifmatchr package (Bioconductor) to perform motif scanning in the sequences of the peak loci, using the threshold of p-value < 5e-5. For both lineage conversion trajectories, we used the chromatin accessibility peak loci with normalized accessibility dispersion greater than 0.5 based on the scATAC-seq data to calculate chromVAR scores for the TFs. We projected chromVAR scores to the force-directed layout of the metacells based on the feature representation from the selected highly variable peak loci as described previously for visualization (**Figure 6B** and **Figure S7A**). Higher chromVAR scores correspond to greater relative accessibility of the genomic regions with potential binding sites of the TF queried.

For the integrated scRNA-seq and scATAC-seq data analysis, chromVAR scores of a TF in each scATAC-seq metacell were projected to the layout of the scATAC-seq metacells in the shared feature space of the scRNA-seq and scATAC-seq metacells. For comparison, gene expressions of the corresponding TF in each scRNA-seq metacell were projected to the layout of the scRNA-seq metacells in the shared feature space.

### Comparison of chromatin accessibility of peak loci across cell groups

We performed differential analysis of chromatin accessibility of selected highly variable peak loci for metacells in the XEN-to-iPS trajectory, to identify peak loci with distinctive accessibility patterns in specific cell groups, or dynamic accessibility patterns across cell groups during the conversion from XEN to iPS cells. There are 7 cell groups (group 1, 2A, 2B, 3, 4A, 4B, and 5) annotated in the XEN-to-iPS trajectory as previously described. We performed two-sample Kolmogorov-Smirnov (K-S) test for each highly variable peak locus to compare the distributions of locus accessibility between a given pair of cell groups, using a threshold of p-value < 1e-4 to identify peak loci that exhibit significant difference in accessibility distributions between the corresponding two groups. More specifically, in **Figure 6C**, we show the chromatin accessibility changes across the different groups of metacells using the identified differential accessible peak loci between group 4B and group 5 of the metacells in the XEN-to-iPS trajectory.

We further identified chromatin accessibility peak loci that were detected in both XEN-to-iPS and ES-to-iXEN trajectories. Given two peak loci from each of the two trajectories, we define them as existing in both trajectories if they overlap with each other. To increase the number of identified peak loci that overlap between the two trajectories, in addition to the originally selected highly variable peak loci, we also use a threshold of normalized accessibility dispersion above 2.0 (*min_disp* parameter in *pp.highly_variable_genes* in the Scanpy package for detection of highly variable peaks as described above) to select an increased set of highly variable peak loci for both trajectories. We then performed differential accessibility analysis for this set of co-existing peak loci between specific pairs of cell groups in a given trajectory. The chromatin accessibilities of the identified set of shared peak loci across cell groups in the ES-to-iXEN trajectory were visualized by heatmap (**Figure 6D**).

### Generating pseudo-bulk ATAC-seq data based on cell groups in the conversion trajectories

Pseudo-bulk ATAC-seq data from the scATAC-seq data were generated based on annotated cell groups in the XEN-to-iPS and ES-to-iXEN trajectories. Each cell group was used to construct a pseudo-bulk sample. In each trajectory, for each cell group and each chromatin accessibility peak locus, we used an average of the chromatin accessibilities of a locus across metacells in the cell group as the accessibility of the locus in the pseudo-bulk ATAC-seq data of the corresponding cell group.

### ChIP-seq analysis

V6.5 ES and IM8A-1 XEN cells were collected in duplicated at ∼25 million each. Cells were crosslinked in 1% PFA in PBS for 10 mins at room temperature and quenched with 125mM glycine for 5 mins at room temperature. Cells were then washed twice with PBS and resuspended in lysis buffer (10mM Tris pH8, 1mM EDTA and 0.5% SDS) at 2×10^7^ cells per 400μl. To shear chromatin, samples were sonicated using a Bioruptor® Pico sonication device (Diagenode) for 12 cycles, 30 seconds on/30 seconds off then pelleted at the maximum speed for 10 mins at 4°C. The supernatant was diluted 5x with dilution buffer (0.01% SDS, 1.1% Triton X-100, 1.2mM EDTA, 16.7mM Tris pH8 and 167mM NaCl), then incubated with primary antibodies at 4°C overnight. Protein G Dynabeads^TM^ (Invitrogen) were blocked at 4°C overnight using 100 ng per 10μl of beads. The next day, beads were added to samples at 20 μl per sample for 3 hrs at 4°C. Using a magnet to stabilize the beads, they were washed twice in low-salt buffer (0.1% SDS, 1% Triton X-100, 2mM EDTA, 150mM NaCl and 20mM Tris pH8), twice in high-salt buffer (0.1% SDS, 1% Triton X-100, 2mM EDTA, 500mM NaCl and 20mM Tris pH8), twice in LiCl buffer (0.25M LiCl, 1% NP-40, 1% deoxycholic acid, 1mM EDTA and 10mM Tris pH8) and once in TE buffer (10mM Tris pH 8, 0.1mM EDTA). Subsequently, the DNA was eluted from the beads by incubating with 150μl elution buffer (100mM NaHCO_3_ and 1% SDS) for 20 mins at 65°C with vortexing using Eppendorf ThermoMixer C (Eppendorf). The supernatant was collected, reverse crosslinked by incubation overnight at 65°C in the presence of proteinase K (Roche), and cleaned by RNase A (Thermo Scientific) treatment for 1 hr at 37°C; the DNA was purified using a DNA clean and concentrate kit (Zymo Research).

Single-end sequenced reads from replicate ES and XEN cells with corresponding input samples were aligned with the use of Bowtie2 (version 2.3.4.1) to mouse genome (mm10) (Langmead and Salzberg, 2012) using “—local-very-sensitive-local” option. Filtering of low quality reads (MAPQ<20), duplicate reads, chrM reads and blacklisted regions was performed with the use of Samtools (Li et al., 2009), “MarkDuplicates” from picard tools and bedtools (Quinlan and Hall, 2010). Filtered reads were used to call both ‘narrow’ and ‘broad’ peaks with MACS2 (version 2.1.1) (Zhang et al., 2008) and default settings with the use of corresponding input for each cell line. Peaks within a distance of a nucleosome were merged into one peak (distance <147 bp) for each replicate. Common peaks between replicates were considered valid and the rest non overlapping peaks were removed from downstream analysis.

### LOLA enrichment analysis

LOLA (version 1.8.0) (Sheffield and Bock, 2016) software in R was used to calculate enrichment of transcription factors and histone modifications in ATAC-seq peaks on mouse genome (mm10). LOLA database was expanded based on available published ChIP-seq data for ES and XEN cells. Enrichment of ChIP-seq experiments was estimated by comparing the enrichment of selected accessible regions (late open, gradual, late close, transient) to an atlas of accessible regions generated by merging ES and XEN ATAC-seq peaks from our experiments. Significant enrichment of transcription factors and histone modifications was scored based on p-value levels (<10^-3^).

## KEY RESOURCES TABLE

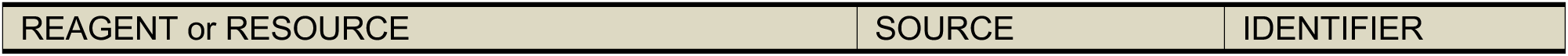
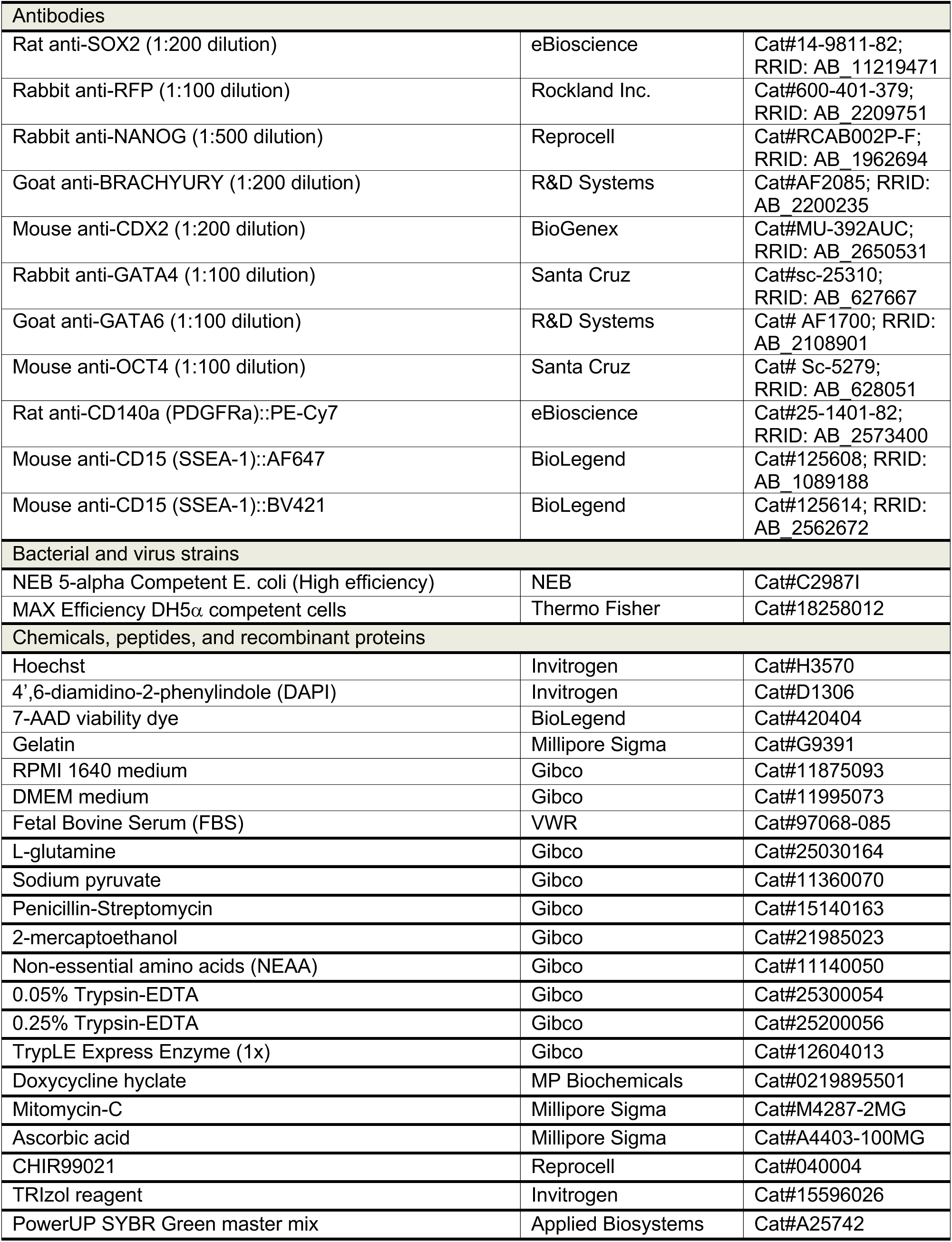
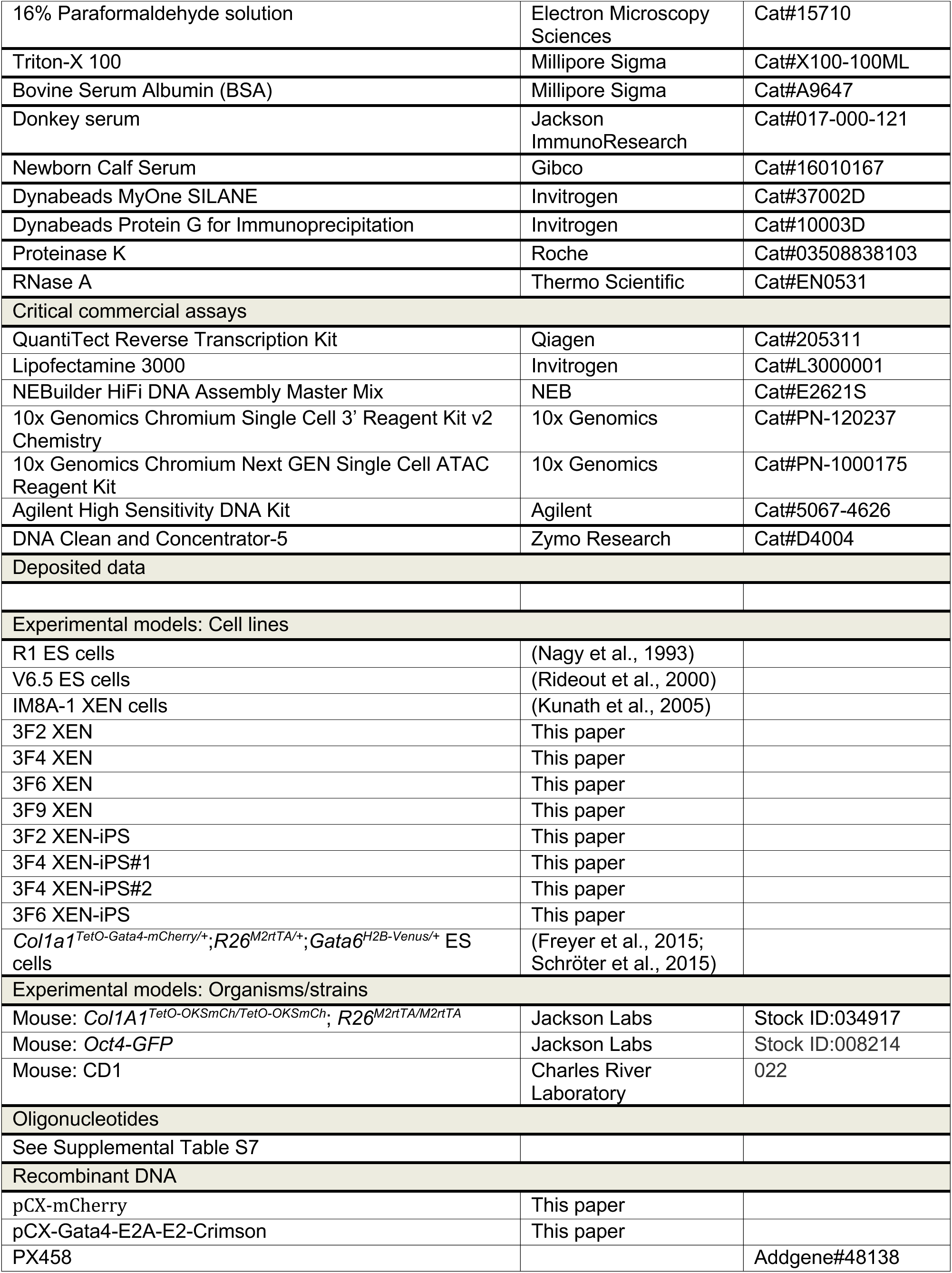
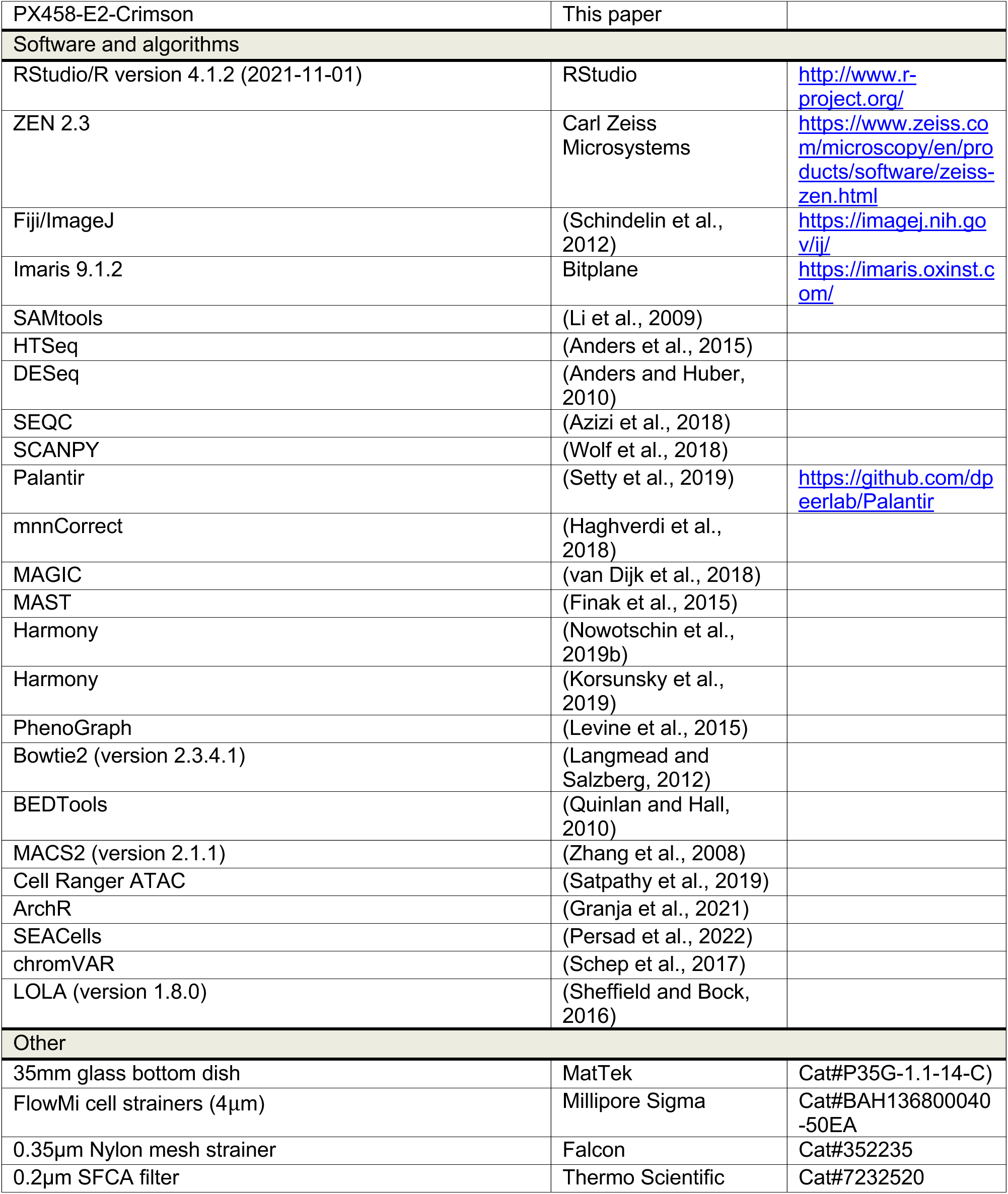

## SUPPLEMENTAL TABLES

**Table S1.** List of differentially expressed genes in terminal states during XEN-to-iPS reprogramming.

**Table S2.** List of GO annotations for differentially expressed genes between intermediate states of XEN-to-iPS and ES-to-iXEN conversion trajectories.

**Table S3.** List of expression trend clusters of genes along XEN-to-iPS conversion pseudotime.

**Table S4.** Highly variable peak loci in scATAC-seq dataset of XEN-to-iPS conversion.

**Table S5.** Highly variable peak loci in scATAC-seq dataset of ES-to-iXEN conversion.

**Table S6.** Peak loci identified as ‘late open’, ‘gradual open’, ‘late close’, and ‘transient open’ from scATAC-seq data fo XEN-to-iPS reprogramming.

**Table S7.** List of oligonucleotides used in this study.

## FIGURE LEGENDS

**Figure S1.**
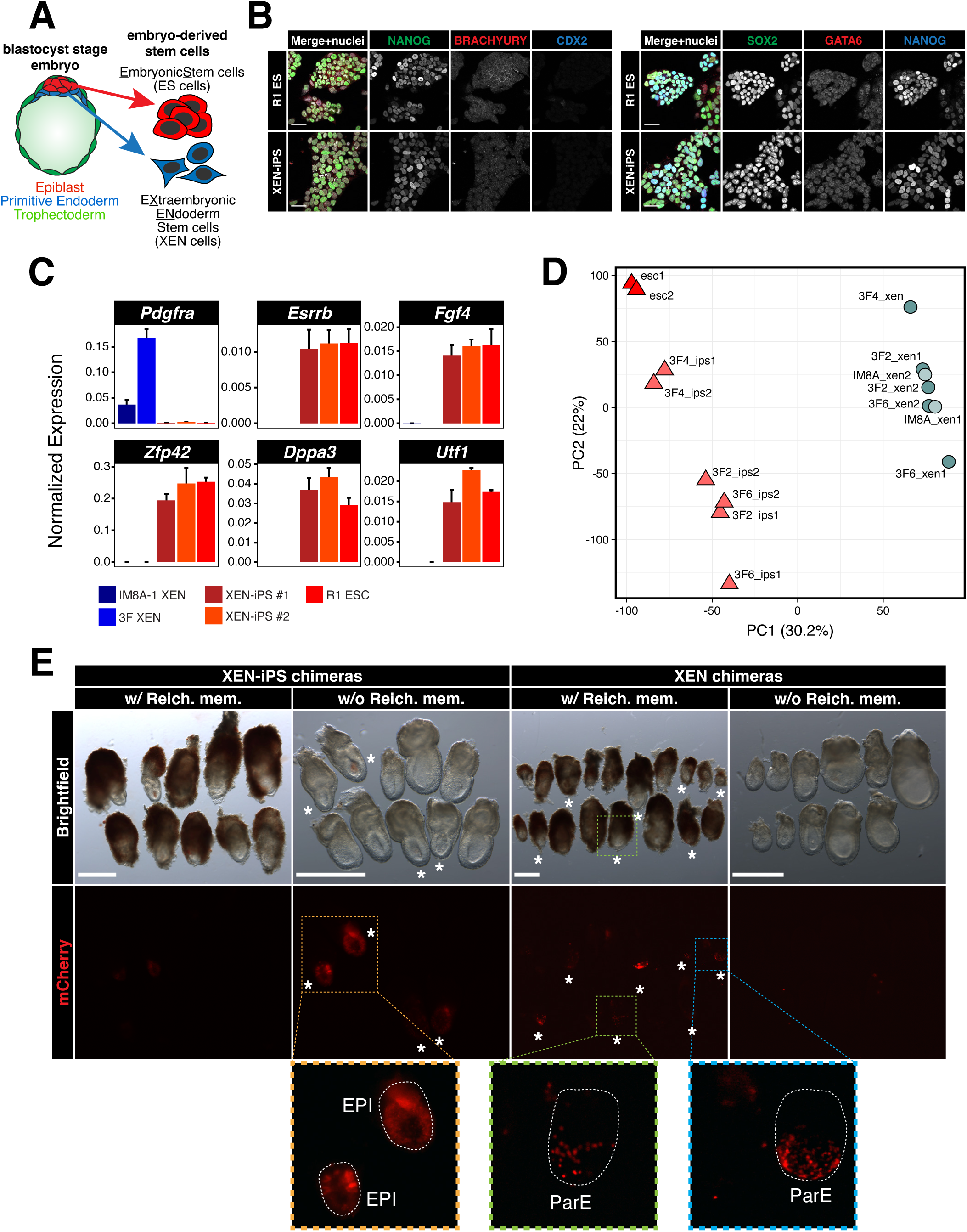
XEN-iPS cells display comparable characteristics and differentiation potential to wild-type ES cells. **(A)** Schematic illustrating the embryo lineage of origin for embryonic stem (ES) cells and extraembryonic endoderm stem (XEN) cells. **(B)** Immunofluorescence staining of wildtype ES cells and XEN-iPS cells with markers of naïve pluripotency, primitive streak, mesoderm and endoderm. Scale bars represent 50µm. **(C)** Gene expression data using RT-qPCR for several XEN and ES cell markers of wildtype XEN (IM8A-1), 3F XEN, two XEN-iPS lines (#1 and #2), and wildtype ES cells (R1). Individual bars show mean expression of three technical replicates normalized to mean expression of two reference genes: *Actb* and *Gapdh*; error bars represent standard deviation. **(D)** Principle component analysis (PCA) plot of bulk RNA-seq data from XEN-iPS cells, wild-type ES cells, and XEN cells. Data was projected onto the top two PCs explaining 30.2% and 22% of the variance respectively. **(E)** Wholemount brightfield and epifluorescence images of XEN-iPS chimeric embryos with and without Reichardt’s membrane demonstrating a lack of contribution of XEN-iPS cells to the ParE- and VE-derived lineages *(left)*, and specific contribution of XEN cells to the ParE lineages *(right)*. Scale bars represent 500µm.

**Figure S2.**
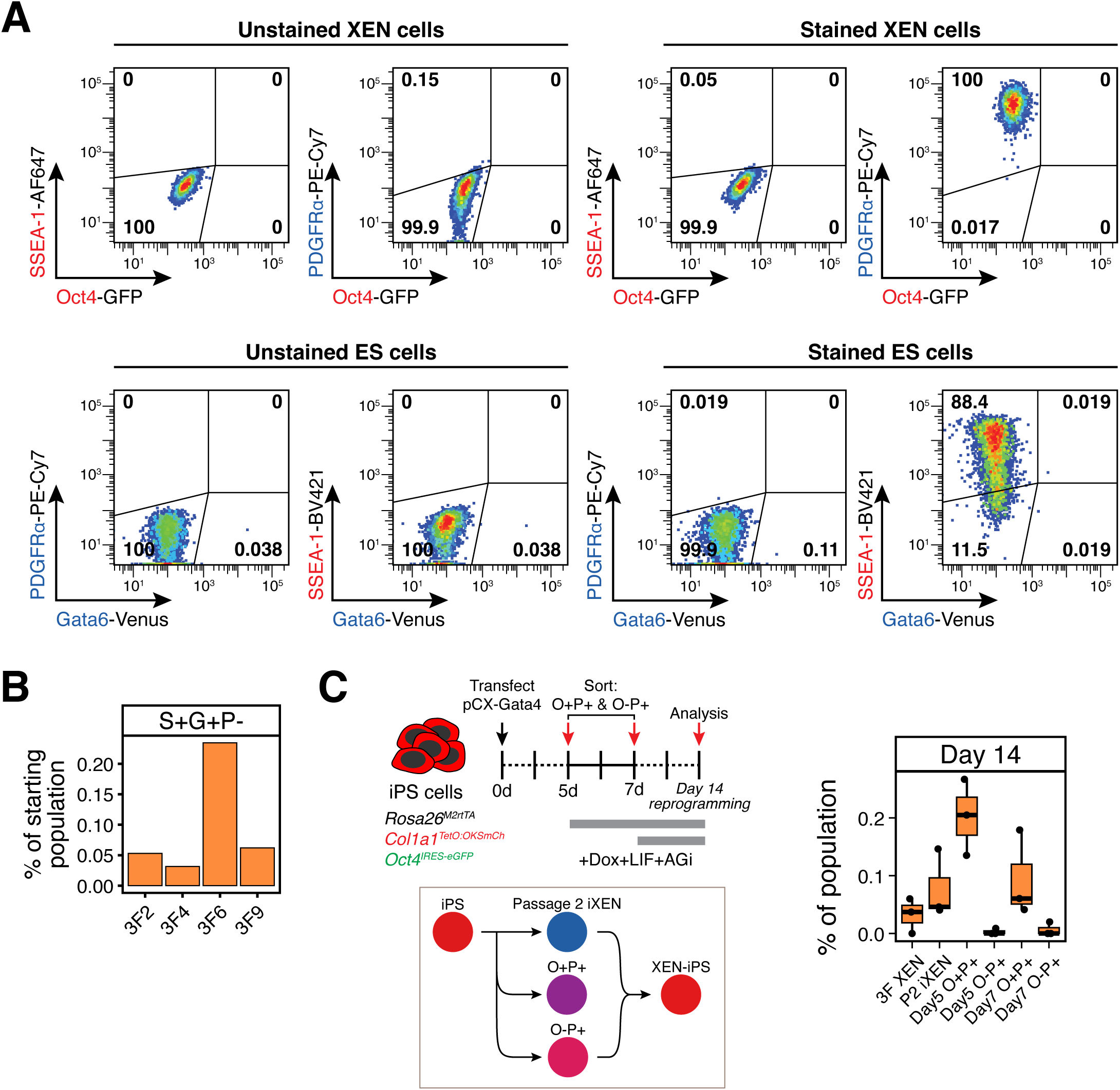
Reciprocal lineage conversions of XEN and ES cells have drastically different kinetics and efficiencies of conversion. **(A)** Gating logic to assess marker expression in live, singlet cells/events. Unstained versus stained XEN or ES cells were used to determine positive and negative gates for each marker. **(B)** Bar chart showing reprogramming efficiency of four different 3F XEN lines. Efficiency was calculated as the percentage of reprogramming cells that are SSEA-1+/*Oct4*-GFP+/PDGFRα-following 28 days of reprogramming. **(C)** *(Left)* Experimental scheme for tracking the reprogramming potential of O+P+ and O-P+ subpopulations sorted at days 5 and 7 following transfection with a Gata4-E2-Crimson expression vector for ES/iPS-to-iXEN conversion. Sorted cells and passage 2 (i.e. nascent) iXEN cells were re-plated in reprogramming conditions for 14 days and resulting percentage of iPS-like cells was determined using flow cytometry. *(Right)* Box plots showing the proportion of the entire population that were S+O+P-at day 14 of reprogramming. Middle line marks the median; lower and upper hinges correspond to the first and third quartiles, respectively. Whiskers extend to 1.5*interquartile range (IQR) from the hinge. Outliers are represented by open circles. N = 3.

**Figure S3.**
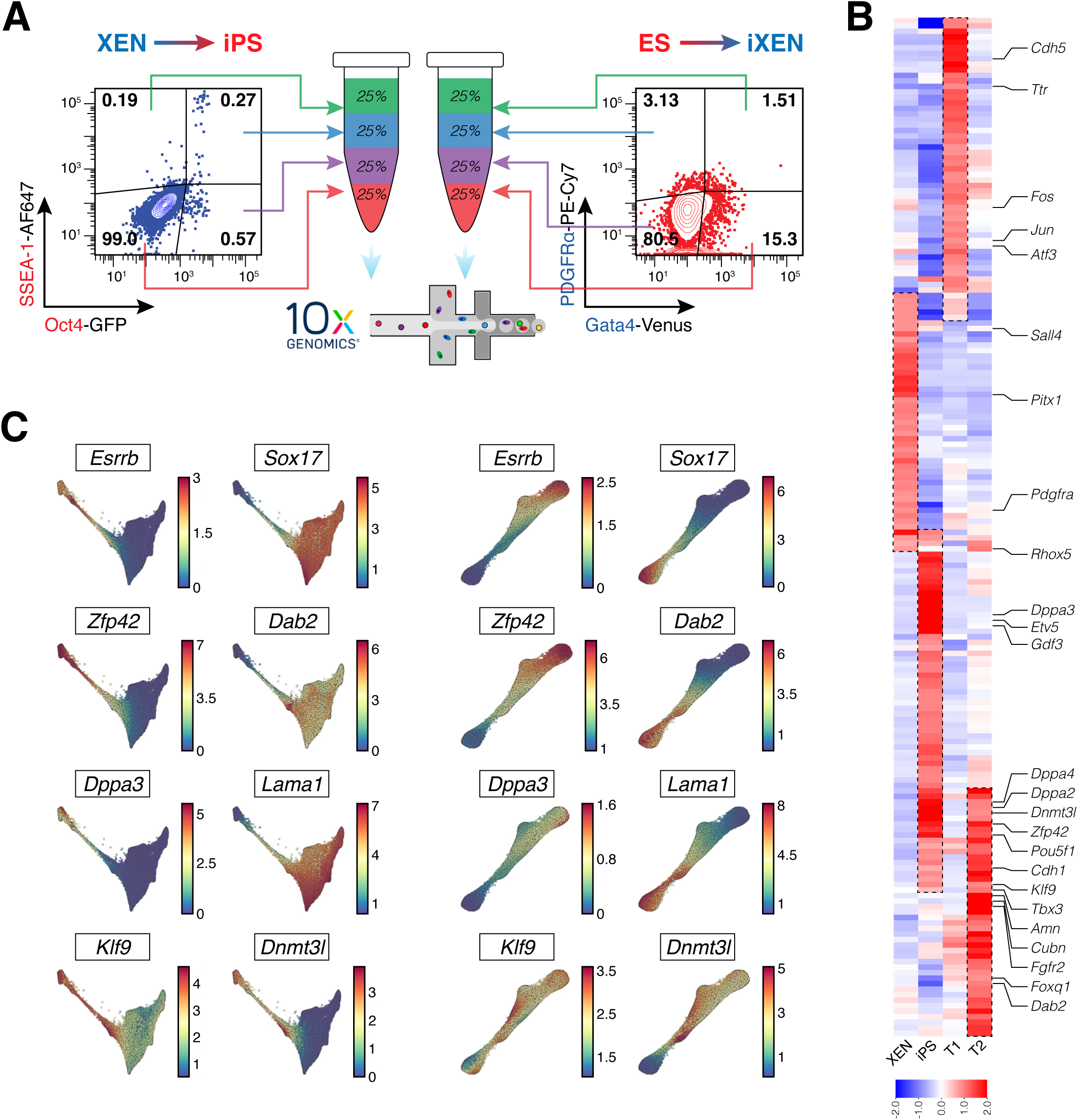
scRNA-seq analyses of XEN-to-iPS and ES-to-iXEN conversions. **(A)** Experimental scheme describing the fluorescence activated cell sorting strategy used prior to encapsulation of Day 7, Day 14 and Day 28 samples of the XEN-to-iPS conversion, and 9h sample of the ES-to-iXEN conversion. Indicated cell populations were reconstituted to represent ∼25% each of the final pool of cells assayed for scRNA-seq. **(B)** Heatmap view of pseudo-bulk differential gene expression between XEN, XEN-iPS, T1 and T2 states. Scale indicates averaged z-score. **(C)** Gene expression patterns of XEN and pluripotency-associated markers. Each cell is colored on the basis of its MAGIC imputed expression level for the indicated gene.

**Figure S4.**
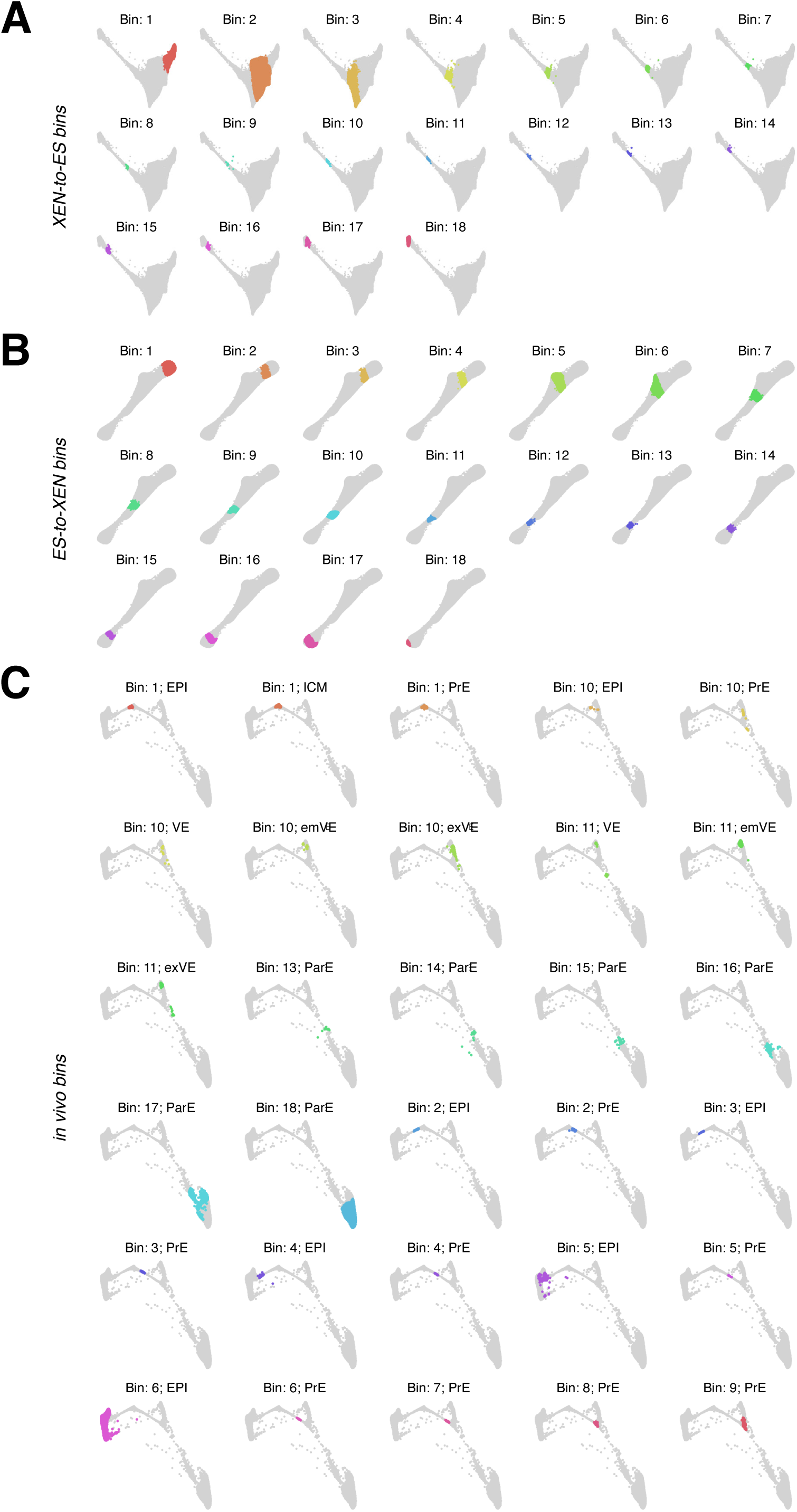
XEN-to-iPS and ES-to-iXEN conversion trajectories follow similar trajectories that approximate *in vivo* cell states. **(A)** Force-directed layout of the XEN-to-iPS trajectory, with individual bins used for the trajectory comparisons. **(B)** Force-directed layout of the ES-to-iXEN trajectory, with individual bins used for the trajectory comparisons. **(C)** Force-directed layout of the *in vivo* trajectory, with individual bins used for the trajectory comparisons.

**Figure S5.**
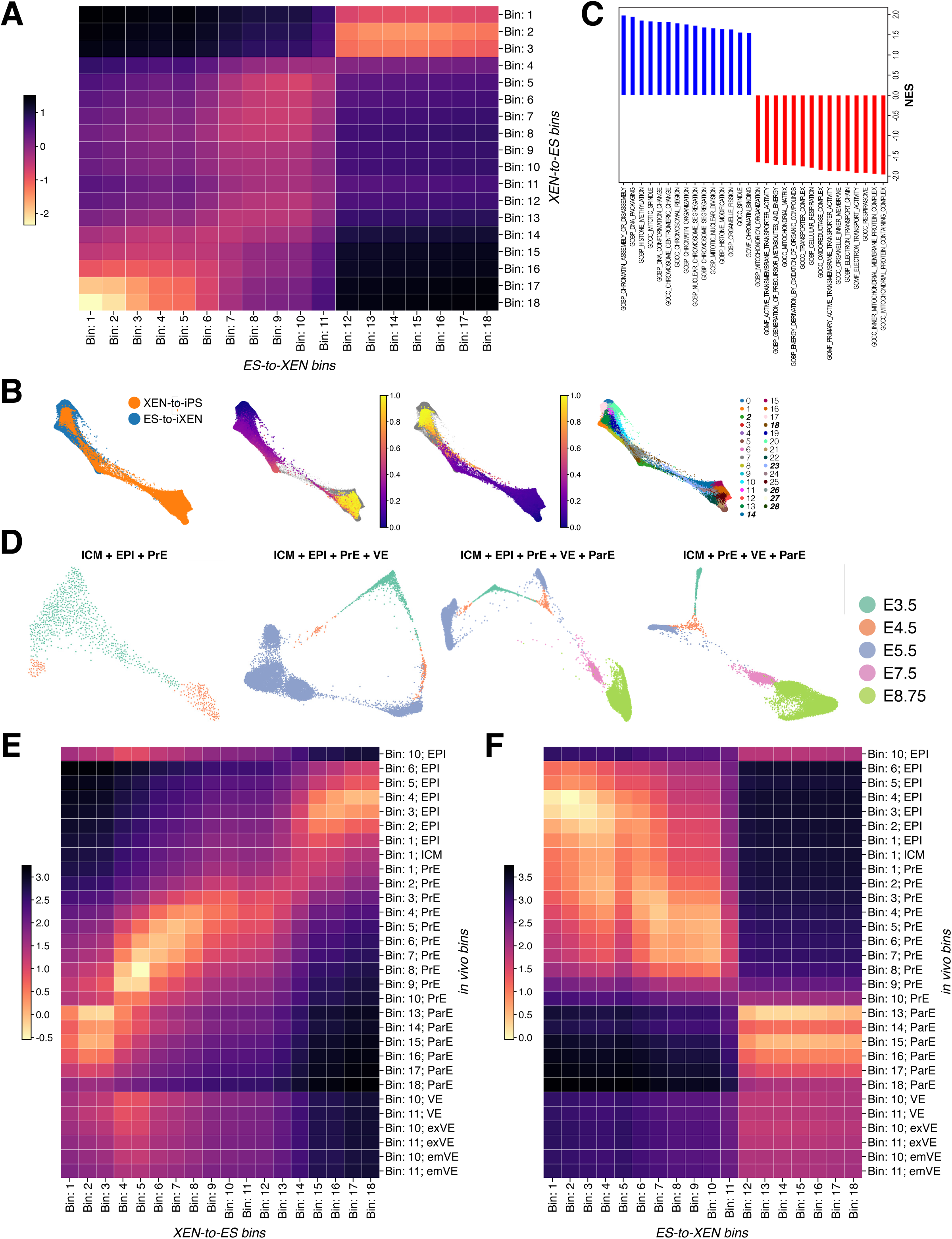
XEN-to-iPS and ES-to-iXEN conversion trajectories follow similar trajectories that approximate *in vivo* cell states. **(A)** Heatmap representation of the relative similarity of different XEN-to-iPS bins to bins from the ES-to-iXEN trajectory. Scale represents relative distance in phenotypic space (lower distance corresponds to increased similarity). **(B)** Force-directed layout of combined XEN-to-iPS and ES-to-iXEN trajectories based on Harmony integration (Korsunsky et al., 2019), colored by (from left to right) trajectory, XEN-to-iPS pseudotime, ES-to-iXEN pseudotime, and PhenoGraph clusters. Intermediate state clusters used to calculate differential gene expression in (C) are bolded and italicized. **(C)** Normalized enrichment scores (NES) of gene ontology pathways calculated for genes upregulated in the ES-to-iXEN trajectory (blue) among intermediate state clusters in (B), or in the XEN-to-iPS trajectory (red). **(D)** Force-directed layouts of combined *in vivo* stages and lineages as labeled above and color-coded as indicated by stage. **(E)** Heatmap representation of the relative similarity of different XEN-to-iPS bins to bins from the *in vivo* trajectory. **(F)** Heatmap representation of the relative similarity of different ES-to-iXEN bins to bins from the *in vivo* trajectory.

**Figure S6.**
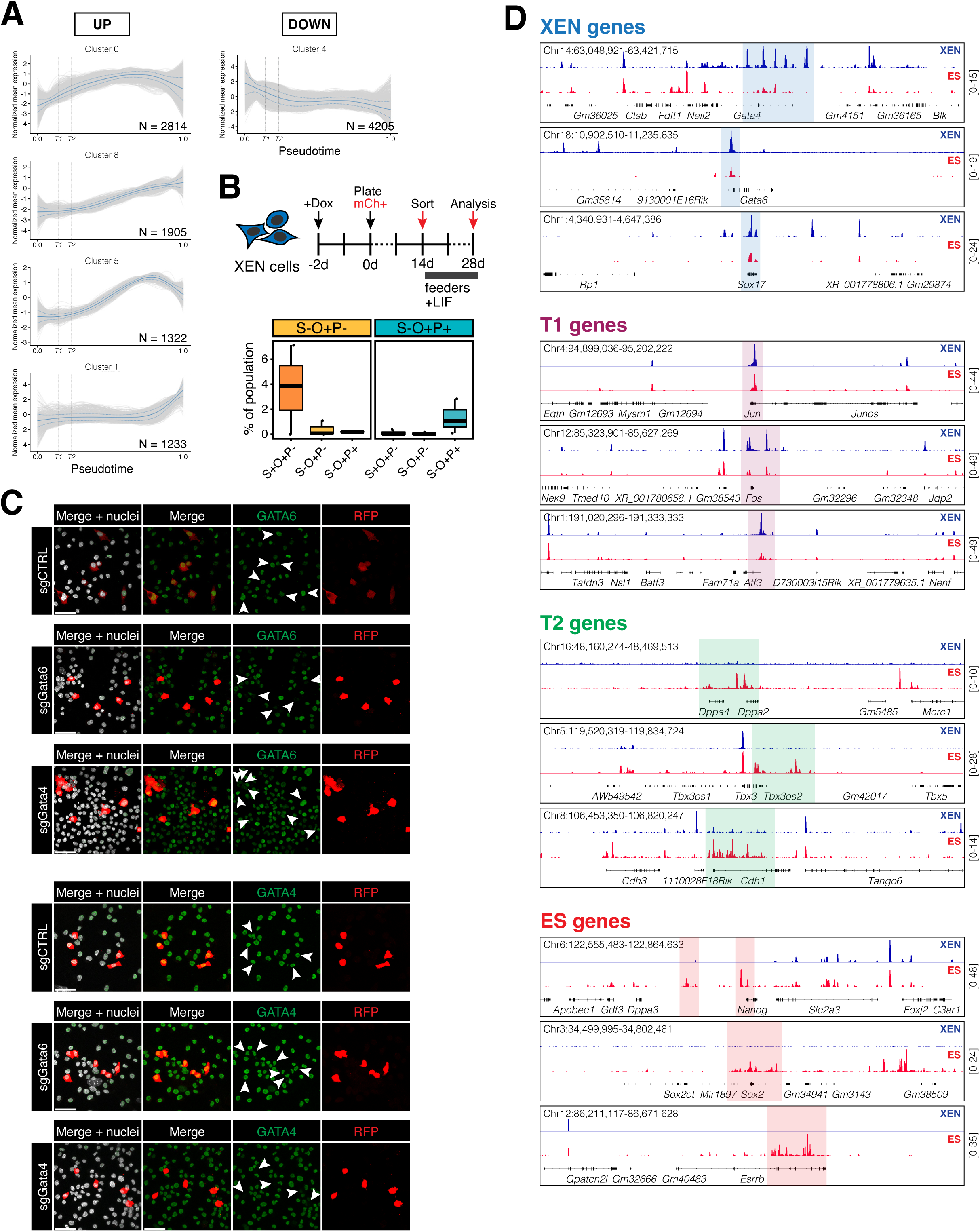
XEN transcriptional network serves as a roadblock to successful XEN-to-iPS reprogramming. **(A)** Gene expression waves over pseudotime of XEN-to-iPS reprogramming. Plots show expression trend of individual genes within each cluster (grey lines). Solid blue line represents mean expression of all genes in the respective cluster. Dotted blue lines represent ± 1 s.d. Vertical dotted lines indicate T1 and T2 terminals states along the pseudotime axis. **(B)** *(Top)* Experimental scheme describing timeline and methodology of tracking reprogramming potential of *Oct4*-GFP expressing subpopulations in the absence of continued transgene expression. S-O+P- and S-O+P+ subpopulations were sorted on day 14 of reprogramming and re-plated in the absence of doxycycline and AGi. Resulting percentage of iPS-like cells was determined after an additional 14 days of culture using flow cytometry. *(Bottom)* Box plots depicting the proportion of the entire population represented by each displayed subpopulation at day 28. Middle line marks the median; lower and upper hinges correspond to the first and third quartiles, respectively. Whiskers extend to 1.5*interquartile range (IQR) from the hinge. Outliers are represented by open circles. N = 3. **(C)** Immunofluorescence staining of XEN cells transfected with Cas9/sgRNA expression vectors targeting *Gata4* or *Gata6*, or non-target control. White arrowheads indicate transfected cells counterstained with anti-RFP antibody. Scale bars represent 50µm. **(D)** Example IGV (Integrative Genomics Viewer) tracks showing ATAC-seq peaks at specific genomic loci surrounding genes representing XEN, ES, T1 and T2 terminal states. Signal values are indicated to the right.

**Figure S7.**
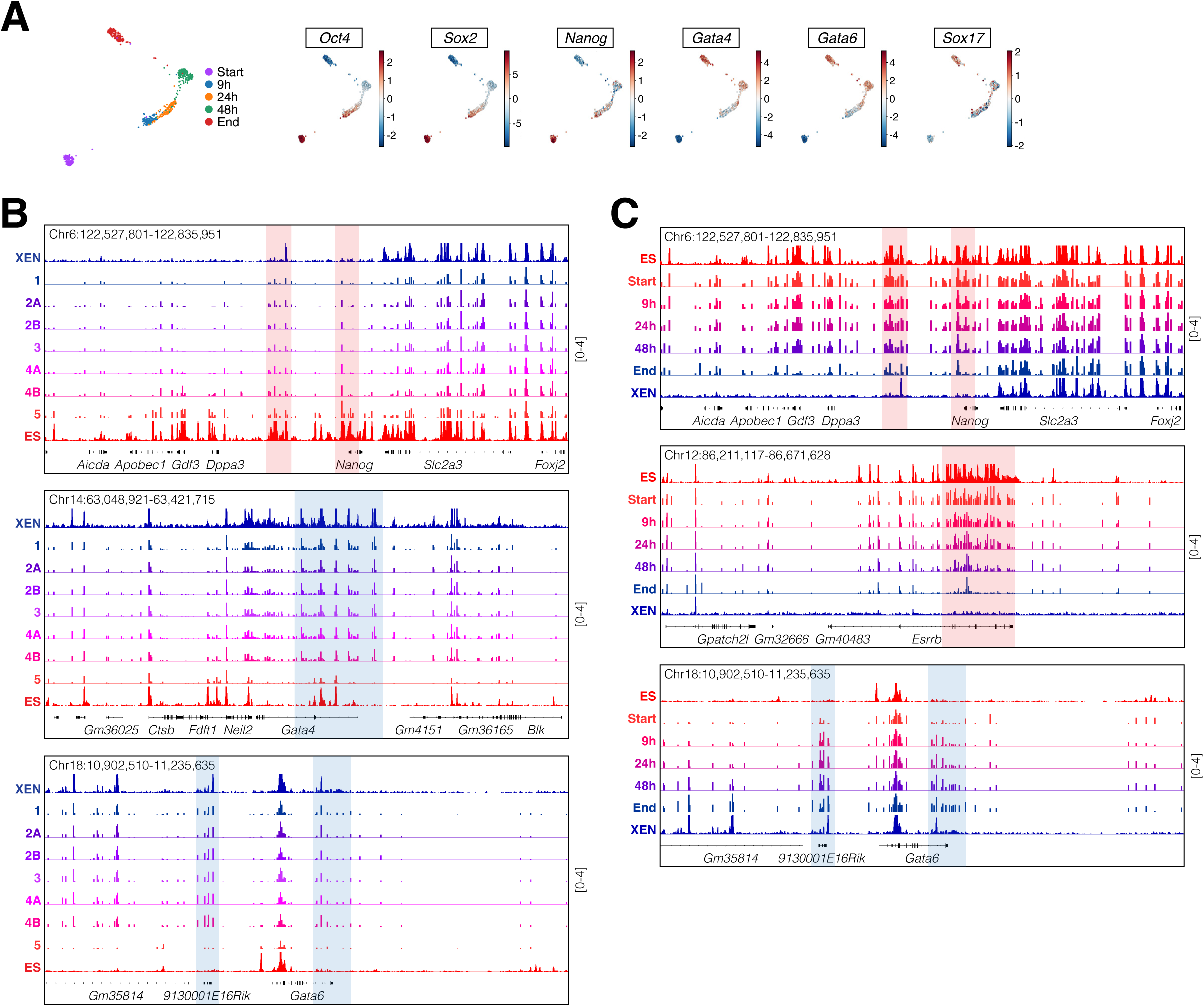
Establishing an EPI-like chromatin state underlies the inefficient conversion of XEN to iPS cells. **(A)** Force-directed layouts showing the combined scATAC-seq dataset for ES-to-iXEN conversion and highlighting the individual timepoints *(left)*, or ChromVAR scores for ES- and XEN-specific TFs *(right)*. **(B)** Example IGV (Integrative Genomics Viewer) tracks showing accessibility peaks of pseudo-bulk scATAC-seq data of XEN-to-iPS conversion. Highlighted are relative accessibility in metacell groups at specific genomic loci surrounding XEN- and ES-specific genes. Signal values are indicated to the right. **(C)** Example IGV (Integrative Genomics Viewer) tracks showing accessibility peaks of pseudo-bulk scATAC-seq data of ES-to-iXEN conversion. Highlighted are relative accessibility at individual timepoints and at specific genomic loci surrounding ES- and XEN-specific genes. Signal values are indicated to the right.

